# ctDNA Release Heterogeneity in Luminal Breast Cancer Revealed by Single-Cell Multi-Omics Spatial Analysis

**DOI:** 10.64898/2026.09.27.754855

**Authors:** Hengyi Xu, Pengming Pu, Binliang Liu, Yuyan Gong, Jiang Wu, Changyuan Guo, Heng Cao, Ziqi Jia, Yuchen Liu, Yansong Huang, Dongxu Ma, Jiayi Li, Zizhao Guo, Tongxuan Shang, Lin Cong, Ruijie Zhou, Quchang Ouyang, Xiang Wang, Yipeng Wang, Jianzhong Su, Yuanyuan Zhang, Jiaqi Liu

## Abstract

**Backgroud:** Breast cancer is a leading cause of cancer-related mortality, and HR^+^/HER2^−^ is the predominant subtype. Molecular pathology, including *PIK3CA* and *ESR1* mutation, informs prognosis and treatment selection. Circulating tumor DNA (ctDNA) offers a non-invasive approach for cancer monitoring, but its clinical utility is limited by tissue–plasma discordance, reflecting heterogeneous ctDNA release. The cellular and spatial determinants of this release remain poorly understood, particularly in HR^+^/HER2^−^ subtype, where hetergeneity across subclones complicates liquid biopsy interpretation.

**Results:** To identify the cellular and spatial determinants of ctDNA release, we developed scMASTER, integrating single-cell transcriptomes and mutations, spatial transcriptomics and matched tumor and plasma DNA sequencing. In ten patients with non-metastatic HR^+^/HER2^−^ breast cancer, eight recurrent malignant epithelial states showed distinct ctDNA release potential. Antigen-presentation, stress, mitosis and mesenchymal-like states showed high release. *PIK3CA* and *ESR1* mutations were linked to distinct functional states and altered ctDNA detectability. Three spatial architectures—EMT-enriched regions, intraductal luminal progenitors and avascular hypoxic zones—showed high release. Tumor–microenvironment interactions were bidirectional: anti-tumoral immunity correlated with reduced release, whereas immunomodulatory components correlated with increased release. A stratification model based on these features identified high-risk, immunosuppressed patients with poor prognosis, validated across external cohorts.

**Conclusions:** Our study establishes a single-cell spatial framework for decoding ctDNA release heterogeneity, linking tumor states, somatic mutations, spatial architecture, and immune interactions to ctDNA release. These findings provide mechanistic insights into tissue– plasma discordance, offer a biological rationale for interpreting ctDNA dynamics, refine patient risk stratification and liquid biopsy-based monitoring in HR^+^/HER2^−^ breast cancer.

## Background

Breast cancer is a leading cause of cancer-related mortality worldwide, and most cases are HR^+^/HER2^−^ [1]. Treatment of HR^+^/HER2^−^ breast cancer has evolved from endocrine monotherapy to molecular pathology-guided precision therapy [2–4]. *PIK3CA* mutations predict benefit from PI3Kα inhibitors combined with endocrine therapy, whereas *ESR1* mutations underlie acquired resistance to aromatase inhibitors and identify candidates for next-generation oral selective estrogen receptor degraders (SERDs) [3–6]. Therefore, longitudinal, comprehensive and tumor-representative molecular profiling is required. Traditional tissue biopsies, however, sample only a limited tumor region at one time point and cannot capture intratumoral heterogeneity [7, 8] or primary-to-metastatic differences in actionable alterations [9].

Circulating tumor DNA (ctDNA) has emerged as a pivotal non-invasive biomarker for liquid biopsy in breast cancer detection[10], treatment monitoring[11], and minimal residual disease assessment [12]. However, its clinical utility as a diagnostic and prognostic tool remains constrained by cancer–ctDNA discordance [13, 14], particularly in non-metastatic disease with lower tumor burden [15]. This discordance arises from heterogeneous ctDNA release across lesions and subclones [7, 8], potentially contributed by cell-state differences, spatial organizations, subclonal mutations and cellular interactions. The mechanisms underlying this heterogeneity remain unclear.

Determining the ctDNA release of different tumor cells is therefore important for relating cancer–ctDNA discordance to tumor biology. However, linking ctDNA to specific cells and cell states remains difficult. Many spatial transcriptomic methods lack true single-cell resolution because of molecular diffusion [16], and high-throughput transcriptomic and mutational measurements are rarely obtained from the same cells [17]. As a result, the cellular origin of ctDNA remains unresolved, limiting mechanistic interpretation and clinical application [18]. Recent advances in single-cell and spatial technologies provide complementary capabilities to address these limitations. Integrating single-cell RNA sequencing (scRNA-seq) with spatial information can localize transcriptional states and cell–cell interactions within tumor niches [19], whereas combining scRNA-seq with mutation data can connect somatic mutations to transcriptional states [20]. Deep learning further enables label transfer and integration across single-cell and spatial modalities [21]. Applied to matched tumor and plasma samples, these approaches can map ctDNA mutations onto defined cell states and spatial architectures, providing a framework to investigate ctDNA release heterogeneity.

Here, we focused on HR^+^/HER2^−^ luminal breast cancer, the subtype in which molecular pathology most directly informs precision treatment, to dissect intra-tumor heterogeneity and track ctDNA. We constructed the single-cell Mutational And Spatial Transcriptomics in Enhanced Resolution (scMASTER) framework by combining full-length RNA sequence transcriptome sequencing (scFAST-seq) [22, 23] with single-cell spatial sequencing (scSpatial-seq) [24] through a deep learning-based method [21]. By integrating single-cell transcriptomes, mutations, spatial data, and liquid biopsy information, we elucidated the mechanisms driving ctDNA release, emphasizing gene expression, functional states, functional mutations, spatial architectures, and microenvironment interactions.

## Methods

### The Overall Analytical Workflow

scMASTER is an integrative analytical framework to summarize our ctDNA tracing workflow. Firstly, scFAST-seq-derived cell type annotations were transferred to scSpatial-seq using SPANN. Afterwards, somatic mutations concurrently detected in tDNA and cfDNA were used for biomarker selection. Patient-specific background dilution was corrected, and genes with significant log₂(cfDNA/tDNA) enrichment across patients were selected, yielding 17 ctDNA-high and 20 ctDNA-low biomarkers. Genes whose expression positively correlated with ctDNA-high mutation rate identified a 77-gene signature. The score was validated by single-cell mutations, the I-SPY2 cohort, and leave-one-out sensitivity analysis.

### Human specimens

A total of ten patients with histologically confirmed HR+ and HER2– breast cancer were enrolled in this study at the Department of Breast Surgical Oncology, Cancer Hospital, Chinese Academy of Medical Sciences (Beijing, China) between 06 2023 and 03 2024. To ensure sufficient ctDNA abundance for reliable downstream analyses while avoiding confounding contributions from distant metastatic niches, we specifically recruited patients with non-metastatic (M0), high-risk tumors characterized by clinicopathological features associated with elevated ctDNA shedding. These features include tumor stage II–III (50% stage II, 50% stage III), mean tumor size >3 cm (range 2.8–5.2 cm), presence of lymphovascular invasion (70%), dermal invasion on microscopy (30%), and a high proliferation index (Ki67 ≥20%, mean 47%). Detailed clinicopathological characteristics for each patient, including age, TNM stage, tumor size, grade, lymphovascular invasion status, microscopical dermal invasion, Ki67 index, and molecular subtype, are provided in Supplementary Table 1.

From each patient, matched specimens were collected: (1) Tumor tissue was obtained from the resected primary tumor and immediately divided along its largest cross-section. One-half was processed for scFAST-seq and targeted tDNA sequencing; the other half was embedded in OCT compound for scSpatial-seq. (2) Peripheral blood (10–15 mL) was drawn into EDTA tubes before surgery. Plasma was separated by double centrifugation (1,600 × g for 10 min, then 16,000 × g for 10 min) and stored at –80 °C until cell-free DNA (cfDNA) extraction. PBMCs were isolated by Ficoll density gradient centrifugation and used as germline controls. All samples were processed immediately after collection. Histopathological evaluation of hematoxylin and eosin (H&E)-stained sections from each tumor was performed by two independent pathologists to confirm diagnosis, assess tumor cellularity, and annotate regions of interest for spatial analysis. Representative whole-slide H&E images are provided in Supplementary Data 1. The study was approved by the Ethics Committee of the Cancer Hospital, Chinese Academy of Medical Sciences (approval number NC2024G-846).

Additionally, we recruited a clinical cohort of 91 late-stage breast cancer patients with ctDNA (the Changsha cohort), data from 200 breast cancer patients with H&E-stained whole-slide images (the Beijing cohort), a published single-cell transcriptome (the Jia-2023 dataset)[25] and a published spatial transcriptome (the Wu-2021 dataset)[26], along with data from two public clinical datasets (TCGA[27] and METABRIC[28]) for external validation (SI Appendix, Table S2 and Methods).

### The Changsha cohort

This study was conducted at the Hunan Cancer Hospital, Central South University, and included patients with advanced breast cancer who underwent ctDNA testing between January 2016 and December 2021. The patient consent form was approved by the Human Research Ethics Committee at the Hunan Cancer Hospital (NO2017YS031), and the protocol has been registered on ClinicalTrials.gov with the number NCT05079074. Each participant provided written informed consent to participate in the study.

### Human sample collection and processing

Primary breast cancer biopsies were obtained from the Department of Breast Surgical Oncology, Chinese Academy of Medical Sciences Cancer Hospital. Tumor tissues were isolated from resected tumors and stored in MACS Tissue Storage Solution-TM at 4 °C. After harvesting, tissues were washed in ice-cold RPMI1640 and dissociated using Multi Tissue Dissociation kit 2 from Miltenyi Biotec according to the manufacturer’s instructions. DNase treatment was optional according to the viscosity of the homogenate. After erythrocytes removal (Miltenyi), cell number and viability were estimated using a Fluorescence Cell Analyzer (Countstar) with AO/PI reagent, and then debris and dead cell depletion (Miltenyi) were determined on live cell results. Finally, fresh cells were washed twice in the RPMI1640 and then resuspended in 1×PBS and 0.04% bovine serum albumin at a concentration of 1×10^6^ cells per mL. Tissue samples were then divided along its largest cross-section; one half underwent scFAST-seq and tDNA sequencing (532 genes; table S1), while the other half was for scSpatial-seq. Blood samples were fractionated to obtain plasma and PBMCs, which were respectively used to extract cfDNA and genomic DNA and underwent DNA sequencing.

### scFAST-seq library preparation and sequencing

Single-cell RNA-Seq libraries were prepared using SeekOne® Single Cell Whole Transcriptome Kit according to the manufacturer’s instructions (SeekGene Catalog No.K00801). Briefly, an appropriate number of cells were mixed with reverse transcription reagents and added to the sample wells of the SeekOne® DD Chip S3 (Chip S3). Then, Barcoded Hydrogel Beads (BHBs) and partitioning oil were dispensed into corresponding wells separately in Chip S3. Subsequently, Cell-containing reverse transcription reagents and BHBs were encapsulated into emulsion droplets using the SeekOne® Digital Droplet System. Immediately following transferring emulsion droplets into PCR tubes, fifteen cycles of annealing (ramping from 8 °C to 42 °C) followed by a 5-minute heat inactivation at 85°C were performed to obtain barcoded cDNA. Next, the barcoded cDNA was purified from broken droplets, and then twice PCR reactions were performed to remove the majority of ribosomal and mitochondrial cDNA. AMPure beads were used to purify cDNA from the post-PCR reaction mixture. Finally, one-fourth volume of cDNA was fragmented, end-repaired, A-tailed, and ligated into the sequencing adaptor. DNA amplified by index PCR contains any part of polyA or non-PolyA RNA as well as Cell Barcode and Unique Molecular Index. The indexed sequencing libraries were purified using AMPure beads and quantified by quantitative PCR (KAPA Biosystems KK4824). The libraries were then sequenced on Illumina NovaSeq 6000 with PE150 read length or DNBSEQ-T7 platform with PE150 read length.

### scFAST-seq data processing

*Fastp* (version 0.23. 1)[29] was used to trim primer sequence and low-quality bases of raw reads and collect the basic statistics. We used the SeekSoul Tools pipeline to process the cleaned reads and generate the transcript expression matrix. Firstly, the cell barcodes and UMI sequences were extracted based on the defined pattern of the localization of the barcode, linker, and UMI within a read. The barcode was corrected with a whitelist. The corrected barcodes, together with UMI, were put in the header of their corresponding reads. Secondly, the reads were mapped to the reference genomes using *STAR* (version 2.5. 1b)[30]. Then, the reads with barcode and UMI information were assigned to transcriptome using featureCounts of package *Subread* (version 1.6.4)[31]. Parameters “ -s ” and “ -t ” of featureCounts vary with different types of chemistries and regions. For the parameter “-s”, “-s 1” was used for products with 3’ chemistry, and “-s 2” was used for products with 5’ chemistry. Another parameter “-t exon” was used for read counting only with exon, and “-t transcript” was used for read counting with exon and intron. we also set the parameter “ -fracOverlap ” to 0.5. Other parameters remain default. Finally, similar to the raw_feature_bc_matrix results of *Cell Ranger* (version 3.1.0)[32], the raw UMI count matrix according to barcodes and transcripts was generated.

A cell-calling algorithm was used to filter the raw UMI count matrix and get the cell-only filtered_feature_bc_matrix. The algorithm was similar to that of Cell Ranger and *EmptyDrops* (version 3.8)[33], which used a cutoff based on the total UMI counts of each barcode to identify cells. This step identified the primary mode of the high RNA content cells. Then the algorithm used the RNA profile of each remaining barcode to determine if it is an “empty” or a cell containing partition. This step captured the low RNA content cells whose total UMI counts might be similar to the empty wells.

### scFAST-seq mutation calling

To identify somatic mutations from scFAST-seq data, we developed a stringent and multi-layer filtering pipeline. Raw sequencing reads were first quality-trimmed using fastp (v0.23.1) with parameters ‘cut_tail_window_size=1’, ‘cut_tail_mean_quality=3’, ‘cut_tail=Truè, ‘unqualified_percent_limit=80’, and ‘length_required=60’. Clean reads were processed with the SeekSoul® Tools mut module, which removes primer sequences, corrects cell barcodes, and aligns reads to the GRCh38 reference genome using STAR (v2.5.1b). Mutation calling was performed with VarScan2 (v2.4.4) in both single-sample and matched tumor-PBMC paired modes. To minimize false positives, we applied the following criteria to each candidate variant: (i) supported by at least 2 unique molecular identifiers (UMIs), 2 cell barcodes, and 2 reads (AD≥2, DP≥2); (ii) variant allele frequency (VAF) ≥0.03; (iii) strand odds ratio (SOR) ≤1 for single-nucleotide variants and ≤10 for indels; (iv) not present at a frequency >1% in the ExAC database (all populations and East Asian) to exclude common germline polymorphisms; (v) each UMI must be supported by at least two PCR duplicate reads with an internal VAF ≥0.9 to eliminate early PCR errors; (vi) the mutation is detected in at least two independent cells from the same sample; (vii) for each cell, the mutant allele fraction (AF) is between 0.2 and 0.8 inclusive. Cells with AF outside this range are excluded due to potential allelic imbalance. Somatic status was determined by requiring the mutation to be absent from the matched PBMC sample (<1% VAF in PBMC) and present in the corresponding tumor DNA (>1% VAF in tDNA). The reliability of mutation detection was confirmed by comparing somatic mutations identified in scFAST-seq with those from ultra-deep targeted sequencing of matched tDNA: 98.8% of scFAST-seq-detected mutations were also found in tDNA, and VAFs showed a strong positive correlation (Pearson *r* = 0.72, *P* < 0.001), with no significant association with gene expression levels (*P* = 0.34), confirming that the mutation calls were not biased by transcriptional abundance. Because mutations were first identified in tumor DNA (which is not subject to RNA editing), this DNA-first approach inherently avoids false positives derived from RNA editing.

### scFAST-seq quality control

To ensure the reliability of single-cell transcriptomic and mutation data, we performed comprehensive QC at multiple stages of the scFAST-seq workflow. Raw sequencing reads were first processed with fastp (v0.23.1) to remove adapter sequences and low-quality bases, yielding high-quality clean reads with an average clean ratio of 96.6% (range 95.8–97.9%) across the ten samples. The base quality was excellent, with mean Q20 and Q30 values of 96.5% and 91.0% (Data S2). Clean reads were aligned to the GRCh38 reference genome using STAR (v2.5.1b). On average, 89.3% of reads mapped to the genome (range 83.5–92.6%), with a high fraction (mean 57.7%) mapping to the middle of gene bodies, confirming effective full-length transcript capture. Ribosomal RNA contamination was minimal (mean rRNA% = 1.1%, mitochondrial rRNA% = 0.9%). Among confidently mapped reads, exonic regions accounted for a substantial proportion (mean 57.5%, range 47.4–67.4%), indicating enrichment for mature transcripts.

Cell calling was performed using the SeekSoul® Tools pipeline, which combines a total UMI cutoff with an EmptyDrops-like algorithm to distinguish genuine cells from empty droplets. This identified a median of 10,600 cells per sample (range 7,406–24,679). To remove low-quality cells, we applied standard Seurat filters: cells with fewer than 300 unique molecular identifiers (UMIs) or with mitochondrial transcript percentage exceeding 10% were excluded. After filtering, a median of 10,430 cells per sample (range 7,245– 24,097) were retained for downstream analysis, with a median of 2,174 genes and 4,402 UMIs detected per cell (Supplementary Data 2). The relationship between mean reads per cell and median genes per cell was assessed by downsampling; sequencing saturation was reached at approximately 30,000 reads per cell, and our achieved mean depth (33,000–53,000 reads per cell) was well above this threshold. Gene body coverage analysis revealed uniform coverage across the full transcript length, with no substantial 3′ or 5′ bias. All QC metrics were summarized in Supplementary Data 2.

### scSpatial-seq spatial barcode preparation

Single-cell nuclei suspension with spatial barcodes was prepared from fresh frozen tissues using the SeekSpace® Single Cell Spatial Transcriptome-seq Kit(K02501-08). Briefly, Fresh frozen tissues were cryo-sectioned to 10-20 μm on a cryostat (Leica) at -20 °C. The tissue regions of interest were then placed on the SeekSpace® Chip, ensuring there were no folds. A finger was placed on the back of the SeekSpace® Chip to melt the tissue. The SeekSpace® Chip was then placed in the SeekSpace® sc-Spatial Chip Holder and incubated on a Thermocycler Adaptor at 37°C for 90 seconds. After 90 seconds, a Space Chamber was placed on the chip, and 150μl of labeling reagent was added without introducing bubbles. Next, the tissue sections were fixed, fluorescence photographed, and homogenized in pre-chilled lysis buffer using a Dounce homogenizer (KIMBLE #D8938). After washing and filtration, the number of nuclei was estimated using a Fluorescence Cell Analyzer (Seekgene#M002B or Countstar® Rigel S2) with AO/PI reagent, before being placed on ice for further use.

### scSpatial-seq Single-cell spatial transcriptome sequencing

Single-cell RNA-Seq library and spatial barcode library were prepared using the SeekSpace® Single Cell Spatial Transcriptome-seq Kit (K02501-08) according to the manufacturer’s instructions. Briefly, the nuclei were evenly divided into 8 PCR tubes, and reverse transcription was carried out on 600-30,000 nuclei in each PCR tube, with a different labeled reverse transcription primer added to each tube. Fifteen cycles of annealing (ramping from 8 °C to 42 °C) were performed to enhance primer hybridization and intracellular reverse transcription efficiency. After reverse transcription, the nuclei were washed twice to remove residual primers and pooled together.

Subsequently, an appropriate number of nuclei were combined with ligation reagents and added to the sample wells of the SeekOne® DD Chip S3 (Chip S3). Barcoded Hydrogel Beads (BHBs) and partitioning oil were then dispensed into the corresponding wells separately in Chip S3. The cell-containing ligation reagents and BHBs were encapsulated into emulsion droplets using the SeekOne® Digital Droplet System. Immediately after transferring the emulsion droplets into PCR tubes, a 60-minute incubation at 20 °C followed by a 10-minute heat inactivation at 65°C was performed to obtain barcoded cDNA and spatial barcodes. The barcoded cDNA and spatial barcodes were then decrosslinked and recovered from cells in droplets. To obtain more Template-Switched cDNA, a second reverse transcription was performed followed by a PCR pre-amplification. The pre-amplified product was used as input for both spatial barcode library construction and cDNA construction. Finally, sample indexes were added to the pre-amplified product during spatial barcode library construction via PCR. After cDNA purification, 20 ng of cDNA was amplified by index PCR. The indexed sequencing libraries were purified using AMPure beads and quantified by quantitative PCR (KAPA Biosystems KK4824). The single-cell RNA-Seq library and spatial barcode library were then sequenced on the Illumina NovaSeq 6000 with PE150 read length or the GeneMind URFSeq 5000 platform with PE150 read length.

### scSpatial-seq quality control

To ensure the reliability of single-cell spatial transcriptomic data, we performed comprehensive quality control at multiple stages of the scSpatial-seq workflow. For each of the ten tumor samples, tissue sections were processed using the SeekSpace® platform, and sequencing libraries were generated as described above. Raw sequencing reads were first assessed for barcode and UMI quality. Across all samples, the median valid barcode rate was 82.3% (range 79.8–85.1%), the median Q30 base quality in barcodes was 89.5% (range 88.2–91.3%), and the median Q30 in UMIs was 96.8% (range 95.9– 97.6%). On average, 93.8% of reads (range 92.1–95.4%) mapped to the reference genome. To evaluate sequencing depth sufficiency, we further performed downsampling analysis. Saturation was reached at approximately 30,000–50,000 mean reads per cell, with a corresponding median of 1,500–2,500 genes detected per cell. Our actual sequencing achieved a median of 14,200 mean reads per cell (range 12,800–16,100). Sequencing saturation averaged 43.5% (range 41.2–46.8%), indicating a balanced trade-off between depth and complexity.

Spatial barcode labeling precision was assessed by analyzing centroid distance distributions. Across all samples, the median radii at the 50th, 75th, and 95th percentiles were 15.3 μm, 25.6 μm, and 51.2 μm, respectively, confirming accurate cell positioning. The median number of cells with detectable spatial barcodes was 38,750 per sample (range 32,100–45,200); after removing contaminants, a median of 21,800 cells (range 18,400–24,900) were retained within tissue coverage areas. For the spatial barcode library, the median valid cell barcode rate was 86.5% (range 84.2–88.7%), and the median valid spatial barcode rate was 72.8% (range 70.5–75.1%). The ratio of valid spatial UMIs averaged 70.1% (range 68.3–72.4%), with a median of 9.2 million valid spatial UMIs per sample (range 8.1–10.5 million). Spatial barcode saturation averaged 79.5% (range 77.0–82.0%), and the median number of spatial UMIs per cell was 52 (range 46–59), providing sufficient resolution for downstream neighborhood analyses. Collectively, these metrics confirm the high quality of the scSpatial-seq data. Detailed per-sample QC statistics are provided in Supplementary Data 3.

### cfDNA extraction and sequencing

Plasma sample collection and cfDNA extraction were performed following kit’s standard protocols. cfDNA underwent quality control for integrity, and concentration was subjected to end-repair and adapter ligation with the Fast Library Prep Kit v2.0(iGeneTech, Beijing, China). 750 ng NGS pre-library was performed hybridization capture with the TargetSeq® Pan-Cancer Panel and TargetSeq One® Hyb & Wash Kit v2.0(iGeneTech, Beijing, China). The libraries underwent PE 150 sequencing on the Illumina NovaSeq 6000, and raw data were filtered with Fastp. Clean reads were aligned to the GRCh38 genome using BWA MEM, and variants were identified and annotated with GATK, Samtools, Varscan, and Annovar software.

### tDNA & PBMC DNA extraction and sequencing

Tissue and PBMC sample collection and gDNA extraction were performed following the kit’s standard protocols. gDNA underwent quality control for integrity and concentration. 200 ng of gDNA was sheared to 150-250 bp fragments using a Biorupter (Diagenode, Belgium). The fragmented gDNA was end-repaired and ligated with Illumina adapters using the Fast Library Prep Kit v2.0 (iGeneTech, Beijing, China) to prepare the NGS pre-library. 750 ng NGS pre-library was performed hybridization capture with the TargetSeq® Pan-Cancer Panel and TargetSeq One® Hyb & Wash Kit v2.0(iGeneTech, Beijing, China). The libraries underwent PE 150 sequencing on the Illumina NovaSeq 6000, and raw data were filtered with Trimmomatic. Clean reads were aligned to the GRCh38 genome using BWA, and variants were identified and annotated with Varscan and Annovar software.

### Calculation of tumor tTMB and blood bTMB

Tumor mutational burden (TMB) was calculated as the number of non-synonymous somatic mutations[34] per megabase (Mb) of the targeted coding region. For tissue TMB (tTMB), we considered somatic mutations identified in tumor DNA (tDNA) sequencing that met the following criteria: (i) variant allele frequency (VAF) ≥5% in tDNA[35] ; (ii) absent in matched PBMC (<1% VAF) to exclude germline polymorphisms[34]; and (iii) supported by ≥5000× sequencing depth. Somatic mutations were called using VarScan2 with paired tumor-PBMC analysis, and only non-synonymous exonic mutations were retained for TMB calculation. For blood TMB (bTMB), we applied additional filters to ensure confident detection in cell-free DNA (cfDNA). Mutations were required to have VAF ≥0.1% in cfDNA, a threshold widely adopted for ultrasensitive ctDNA detection[36, 37]. As demonstrated in previous validation studies, targeted sequencing at depths exceeding 5,000× enables reliable variant calling at this allele frequency with high sensitivity (>99% for point mutations) and specificity[36]. TMB values were calculated by dividing the total number of qualifying somatic mutations by the panel size (2.126 Mb, covering 641 genes). The final tTMB and bTMB values for each patient are reported in Supplementary Table 2.

### Construction of the scMASTER framework

To dissect ctDNA release heterogeneity, we developed an integrated analytical pipeline that combines single-cell mutation, transcriptome, spatial, and liquid biopsy data, which we referred to as **scMASTER** (single-cell Mutational and Spatial Transcriptomics in Enhanced Resolution). It is important to note that scMASTER is not a standalone software package, but an analytic framework that integrates multiple existing computational tools and analytical modules. The framework comprises three core components:

1. **SPANN-based label transfer.** To achieve single-cell resolution spatial annotation, we employed SPANN (version 0.0.5)[21], a deep learning method that integrates scRNA-seq reference data with spatial transcriptomic profiles. Spatial data were initially stratified into immune, epithelial, and stromal subpopulations. Subsequently, refined subtyping was performed within each subpopulation. SPANN first uses a pair of interconnected variational autoencoders that project both reference and spatial datasets into a unified latent space. To leverage spatial proximity, a neighbor loss function was incorporated, assigning similar labels to physically adjacent cells. Label transfer was performed using the overlapping set of the top 5,000 most variable genes in scFAST-seq and scSpatial-seq. Model training was executed with parameters: learning rate = 2 × 10⁻⁴, resolution = 0.5, λ_spa = 0.001, λ_cd = 0.001, λ_nb = 10, max iterations = 5,000, with intermediate checkpoints at 2,000 and 4,000 iterations. The accuracy of transfer was validated on non-malignant cells with well-established markers. Comparison of SPANN-assigned labels against manual annotation yielded >90% diagonal concordance in the confusion matrix and F1 scores >0.65 for all cell types, with UMAP visualizations confirming highly similar distributions. For malignant epithelial cells, we confirmed that state-specific gene markers and functional scores derived from scFAST-seq were faithfully preserved after transfer to scSpatial-seq. We further performed permutation-based validation (1,000 random label shuffles). The true state-specific transferred scores were significantly higher than permuted scores (*P.adj* < 0.001 for all states), confirming accurate label transfer.
2. **ctDNA biomarker selection.** To identify mutations that could serve as reliable tracers of ctDNA release, we first established a set of somatic mutations concurrently detectable in scFAST-seq, tDNA, and cfDNA. All mutation types (including synonymous and UTR variants) were retained to preserve traceable markers. Mutations were required to meet the following criteria: cfDNA VAF ≥0.1%, tDNA VAF ≥1%, and PBMC VAF <1%. For each patient, we calculated the patient-specific background dilution rate using these mutations, defined as the mean log₂(mean cfDNA VAF/mean tDNA VAF). For every gene, we then computed its patient-level log₂ fold change (logFC) as log₂(cfDNA VAF / tDNA VAF) and subtracted the background dilution logFC to obtain a corrected logFC. Genes with a corrected logFC significantly different from zero across the ten patients (one-sample t-test, P.adj < 0.05) and with an absolute mean logFC >0.5 were considered candidate biomarkers. Additionally, we required that the direction of change (enriched or depleted in cfDNA) be consistent in at least 50% of patients. To further validate the statistical robustness of each candidate, we constructed 2×2 contingency tables based on read counts and performed Fisher’s exact test for each patient; all genes had *P*.adj < 0.05. This procedure yielded 17 genes significantly enriched in cfDNA (ctDNA-high) and 20 genes significantly depleted (ctDNA-low). Gene Ontology enrichment analysis revealed that ctDNA-high genes were primarily involved in cell growth, apoptosis, and DNA repair, whereas ctDNA-low genes were enriched in immune and normal pathways.
3. **ctDNA release score identification.** To quantify the ctDNA release potential of individual cells based on their transcriptomic profiles, we developed a ctDNA release score. Using scFAST-seq data, we first identified cells that harbored mutations in the ctDNA-high biomarker genes. Only cells with at least 10 reads for the relevant gene panel were included to minimize sampling noise. For each of the 17 ctDNA-high genes, we calculated the per-cell mutation rate (presence/absence) and performed a Pearson correlation analysis between the mutation rate and the expression levels of all genes in epithelial cells. Genes whose expression showed a significant positive correlation with ctDNA-high mutation rate (Pearson *r* > 0.15, *P*.adj < 0.05) were selected to constitute the ctDNA release signature. This procedure yielded a 77-gene set. The ctDNA release score for any given cell was then computed as the average expression level of these 77 genes using the *AddModuleScore* function in *Seurat*.

### Validation of the ctDNA release score

To validate the robustness and biological validity of the ctDNA release score, we performed three independent layers of validation:

1. **Internal validation using scFAST-seq.** We used mutations coexisting in scFAST-seq and tDNA for internal validation (PBMC VAF <1%, tDNA VAF ≥1%). Critically, the mutations used for validation were not in the 37-gene biomarkers or 77-gene ctDNA release signature, excluding technical circularity. For each somatic mutation, we identified cells that harbored “releasing-type” mutations (cfDNA at VAF ≥0.1%) and “non-releasing” mutations. Only cells with ≥10 covered reads at the relevant locus were included. We identified 12 genes that harbored both releasing and non-releasing mutations across different cells with sufficient coverage. For each of these genes, we compared ctDNA release scores between cells carrying the releasing versus non-releasing mutations using the Wilcoxon rank-sum test (Supplementary Table 4). Additionally, we compared ctDNA release scores between cells harboring at least one releasing-type mutation and cells harboring no releasing mutations, obtaining a positive result with *P* < 0.05 by the Wilcoxon signed-rank test.
2. **External validation in the I-SPY2 cohort.** We performed external validation using the baseline data of I-SPY2 cohort[38], a multicenter, adaptively randomized phase II trial of high-risk breast cancer, with available bulk tumor RNA-seq data and binary ctDNA status (positive/negative). We computed the ctDNA release score by applying the 77-gene signature using normalized average expression and compared scores between ctDNA-positive and ctDNA-negative patients. ctDNA-positive patients exhibited significantly higher ctDNA release scores than ctDNA-negative patients (mean 0.315 vs. 0.189, Wilcoxon test *P* = 0.045). Moreover, the ctDNA release score correlated positively with proliferation, stress response, and apoptosis pathways (Pearson *r* > 0.3, *P* < 0.01 for each). This external validation confirms that the ctDNA release score is generalizable beyond our single-center discovery cohort.
3. **Leave-one-out validation for patient bias.** To ensure that our findings were not driven by these outlier patients, we performed leave-one-out sensitivity analyses. We first re-ran the biomarker selection pipeline. When the two high-contributing patients (LuBC04 and LuBC08) were removed simultaneously, 77.8% (28/36) of the biomarkers were retained. The logFC values of biomarker candidates from the full and the reduced cohort showed strong correlation (Pearson *r* = 0.81, *P* < 0.001). We reconstructed the 77-gene ctDNA release score after iterative exclusion of each patient. Across all ten iterations, the leave-one-out scores showed high correlation with the original score (mean Pearson *r* = 0.88, range 0.84–0.92). The associations between the score and proliferation, stress, and apoptosis pathways were preserved in every iteration (all *P* < 0.05). Additionally, the hazard ratio for overall survival in the METABRIC cohort varied by less than 15% across iterations, and the log-rank test remained significant in all cases (*P* < 0.05), confirming the prognostic value of the ctDNA release score.

### Clustering and visualization

The clustering and visualization were finished by *Seurat* (version 4.0.3)[39] with the following steps:

1. Data normalization. LogNormalize, a global-scaling normalization method, was employed to normalize the expression. The expression measurement of one transcript was divided by those of all the transcripts of the cell and multiplied by a scale factor (10,000 by default), and then the result was logarithmically transformed.
2. Detection of highly variable features. FindVariableFeatures was used to get 2,000 features per dataset.
3. Scaling. A linear transformation (’scaling’), a standard pre-processing step prior to dimensional reduction techniques, was applied.
4. Dimensional reduction. PCA on the scaled data was performed, and the first 15 principal components were used in the following steps.
5. Clustering. A graph-based approach was applied to cluster the cells[40].
6. tSNE/UMAP. The non-linear dimensional reduction technique was used to visualize and explore these datasets.
7. Cluster markers. FindAllMarkers with the default parameters except “logfc.threshold=1” was used to find markers that determined the cell clusters via the differential expression, and the top markers were visualized.

### Meta-cluster-based malignant epithelial analysis

The top 50 preferentially expressed genes from each subgroup were selected to constitute the expression module for the respective meta-cluster. The 34 modules were then aggregated into various recurrent expression programs through hierarchical clustering aligned with their expression profiles. Eight distinct expression programs, indicative of different functions and cellular states, were identified, with cells from clusters within the same program categorized as program cells. The most activated pathways within these program cells were selected by comparative analysis with other cells to elucidate their functional roles.

### Validation of eight-state epithelial subtyping framework

To ensure that the eight epithelial states identified in this study represent robust and generalizable biological programs rather than artifacts of patient-specific heterogeneity, we performed comprehensive validations. We first quantified the patient contribution to each epithelial state. Five states (AP, Gland, Mes, ECM, and ERresp) exhibited single-patient dominance, with one patient contributing >30% of cells within the state (range 53.8–79.8%). To further assess the stability of state-defining gene signatures, we performed leave-one-patient-out analysis for each state. For every state, we iteratively removed the dominant patient and recalculated the top 50 marker genes (FindMarkers function in Seurat) using the remaining patients. The Jaccard similarity index was computed between the original full-cohort marker set and the leave-one-out marker set for each iteration. States with single-patient dominance showed variable stability: AP, Gland, ERresp, and ECM exhibited Jaccard indices below 0.5, while Mes, Mitosis, Stress, and OP maintained Jaccard indices above 0.6, indicating high stability. Then, for the four states with Jaccard indices below 0.5, we stratified cells into two groups: those from the dominant patient (“Main”) and those from all other patients (“Others”). Within each state, we performed AddModuleScore comparing Main versus Others to identify differentially enriched pathways. For all five states, the top enriched pathways were highly concordant between the two groups. Key marker genes for each state also showed consistent expression patterns between Main and Others, further supporting the biological coherence of the state definitions.

To further test the generalizability of our eight-state framework, we integrated several datasets, constructing a single-cell RNA-seq profile comprising 50 breast cancer patients (50BC [37,38]), encompassing diverse molecular subtypes and clinical stages. Raw count data were obtained and processed using the same normalization and scaling procedures as our discovery cohort. Cell clusters were assigned to the state according to their expressions in our pre-defined marker panels. Across the 50BC cohort, 93% of malignant epithelial cells were successfully assigned to one of the eight states. Each state was supported by cells from multiple patients (6–34 patients per state), with a minimum of 50 cells contributed per patient. For each state, the marker genes and functional pathway expression patterns were similar to those of our cohort. These results demonstrate that the eight epithelial states represent recurrent and biologically meaningful programs that are reproducible across independent patient populations and experimental platforms.

### Mutational label transfer on single gene from scFAST-seq to scSpatial-seq

The scSpatial-seq data was not whole-length and lacked single-cell mutation information. Therefore, it was necessary to perform mutational label transfer using scFAST-seq. First, in scFAST-seq, we conducted differential expression analysis between mutated cells and non-mutated cells for specific mutations. A *P*.adj<0.05 and a log2 (fold change)>0.5 were used to obtain the transcriptomic score for the mutation. Subsequently, the mutation scores were transferred to the scSpatial-seq epithelial cells of the same patient using the AddModuleScore function. The mutated cells were identified based on the same mutation ratio in the epithelial group of scFAST-seq.

### External validation of ESR1 mutation signatures

To independently validate the transcriptional signature of the *ESR1* D538G mutation and its association with ctDNA release, we leveraged publicly available bulk RNA-sequencing data from breast cancer cell lines MCF-7[41]. The dataset includes *ESR1* D538G, *ESR1* Y537S, or wild-type control. Counts were normalized to transcripts per million (TPM) for downstream analysis. The 200-gene *ESR1* mutation score was identified in our scFAST-seq data by comparing *ESR1* D538G-mutant cells versus wild-type cells (|log₂FC| > 0.5, *P*.adj < 0.05; Supplementary Table 5). The ctDNA release score was calculated using the 77-gene signature established in this study. Both scores were computed as the average expression after z-score normalization. Cell lines harboring *ESR1* D538G showed significantly higher *ESR1* mutation scores and lower ctDNA release scores compared to wild-type controls, accompanied by elevated expression of mammary differentiation and estrogen response pathways (all Wilcox test *P* < 0.01).

### Spatial regional sequencing

To validate the spatial distribution of somatic mutations and their association with ctDNA release, we performed targeted DNA sequencing on spatially dissected regions from tissue sections immediately adjacent to the scSpatial-seq slices. Serial sections were cut at a depth of 80 μm from the same tissue block, and each section was divided into regions according to the dissection scheme illustrated in Supplementary Data 4. The number of dissected regions per section was adjusted based on tissue area: sections with smaller areas were divided into two regions to ensure sufficient DNA yield, while larger sections were divided into four regions. Genomic DNA was extracted from each dissected region, and all samples passed quality control with total DNA exceeding 300 ng. Targeted sequencing was performed using the same panel as used for tDNA sequencing in the study (641 genes, 2.126 Mbp). The mean sequencing depth across all regional samples was 4,507×, with >99.9% coverage. Somatic mutations were defined by integrating the regional sequencing data with previously established tDNA and PBMC profiles. Only mutations present in tDNA with VAF >1% and absent in PBMC (VAF <1%) were selected for the following analysis. For each patient, mutations were classified into two categories based on their spatial distribution across regions. Spatial-unique mutations were defined as mutations detected exclusively in a single region, while spatial-common mutations were defined as mutations present in all regions examined.

### Analysis of the relationship between ctDNA read count and single-gene expression

The analysis of the relationship between ctDNA release and single-gene expression was divided into two levels: single-cell level analysis using scFAST-seq data and patient-level analysis using an external clinical cohort.

1. At the single-cell level, we calculated the average expression level of each gene in epithelial cells and obtained the average mutation rate in tDNA and the average number of mutated reads in ctDNA for each gene. Subsequently, we performed a correlation analysis between the average epithelial expression levels of all genes and the ctDNA mutation reads normalized by the tDNA mutation rate.
2. In the external cohort analysis, we first identified high and low-expressed genes of all, epithelial, and non-epithelial cells in scFAST-seq data by defining genes with average expression in the top 1/3 as high-expressed and those in the bottom 1/3 as low-expressed. Then, we compared the mutation rates of high and low-expressed genes in the external ctDNA cohort.

### Pathway enrichment analysis

1. Gene Ontology (GO) enrichment analysis of the marker genes was implemented by the *ClusterProfiler* R package (version 4.10.0)[42], in which gene length bias was corrected. GO terms[43] with *P*.adj less than 0.05 were considered significantly enriched.
2. KEGG is a dataset resource for understanding high-level functions and utilities of the biological system, such as the cell, the organism, and the ecosystem, from molecular-level information, especially large-scale molecular datasets generated by genome sequencing and other high-throughput experimental technologies (http://www.genome.jp/kegg/)[44]. We used the clusterProfiler R package to test the statistical enrichment with *P*.adj less than 0.05 of marker genes in KEGG pathways.
3. The Reactome Knowledgebase (https://reactome.org) provides molecular details of signal transduction, transport, DNA replication, metabolism, and other cellular processes as an ordered network of molecular transformations—an extended version of a classic metabolic map, in a single consistent data model. We used the clusterProfiler R package to test the statistical enrichment with *P*.adj less than 0.05 of marker genes in Reactome pathways.

### Module score analysis based on Msigdb dataset

As different breast cancer subtypes have different characteristics, functional comparison is executed on the level of subtypes. In hypoxia analysis, subgroups representing the hypoxia feature were selected based on five hypoxia scores from Msigdb dataset[45]. Functional scores were calculated using the AddModuleScore function in *Seurat*. Scores were centered by subtracting the average value of all tumor samples. Subgroups with the highest hypoxia score were specified as hypoxia subgroups.

### Copy number variation inference

CNV signals for each chromosome region of each malignant cell were determined using the hidden Markov model in *infercnv* package (version 1.6.0)[46]. The “subcluster” mode was used to determine clonal CNV changes. GRCh38 cytoband information is used to convert each CNV to p or q arm horizontal variations for simplification based on its position. Chromosome arms with CNV proportion higher than 40% were annotated as gain or loss. For visualization, *Uphyloplot2* (version 2.3) is used for the autonomic construction of intra-patient phylogenetic tree[47]. GRCh38 gene position information was used to calculate the copy number variation of several tumor driver genes.

### Differential mutation analysis and differential mutation score

Utilizing Fisher’s exact test, mutation rates at a specific locus were compared between two populations. If the *P*.adj from Fisher’s exact test was less than 0.05, the mutation was considered to have significantly increased or decreased in frequency within that population. For example, the differential mutation sites for a specific epithelial subgroup were defined as positive enrichment with mutation sites with a *P*.adj <0.05, comparing this subgroup with others.

### Spatial adjacent analysis

Epithelial state adjacency analysis was divided into global adjacency analysis and single-patient epithelial partition analysis.

1. In the global adjacency analysis, we calculated the average cell distance between any two epithelial subgroups for each patient and then averaged these distances across all patients. However, the resulting matrix was a distance matrix. To further process this, we normalized the data on a column-by-column basis using the formula: x_adjacent_=Normalize(x−x_min_) to obtain an adjacency matrix for each state relative to other states, which was then visualized as a heatmap.
2. In the single-patient epithelial partition analysis, we divided the scSpatial-seq slice of each patient into a 20x20 grid and calculated the proportion of the eight epithelial states within each grid. These proportions were used as epithelial proportion information for spot hierarchical clustering to obtain spatial epithelial partitioning. The choice of K (number of clusters) was based on the gap statistic, following the formula: K = which.max(fgap(K)) Here, the number of clusters corresponding to the maximum gap value was selected as the optimal number.

### Spatial EMT area identification and functional analysis

In the identification of spatial EMT areas, we first quantified the number of Mes state epithelial cells surrounding each cell and generated density plots for each patient. Subsequently, based on the shape of the density plots, if the plot was bimodal, the threshold was set as the average of the two peaks; if the plot was unimodal, the threshold was set at two standard deviations to the right of the peak. After the identification of EMT areas, comparisons of transcriptomic profiles and immune infiltration between different regions could be conducted. Differential expression analysis between cells inside and outside regions yielded a 90-gene spatial EMT score (*FindMarkers* function, *P*.adj<0.05, logFC>0.5). We used Wu-2021, a dataset of spot-based spatial transcriptome of a luminal breast cancer patient, for external validation of this score[26].

### Clinical relevance of EMT-associated ctDNA release score

The external cohort utilized in this analysis comprised 91 patients with late-stage (III/IV phase) breast cancer, each with mutation profiles for both cfDNA and corresponding tDNA, allowing for survival analysis based on mutations. The EMT-associated feature was defined as the union of spatial EMT score (*FindMarkers* function, *P*.adj<0.05, logFC>0.25) and Mes epithelial markers (*FindMarkers* function, *P*.adj<0.05, logFC>1). A patient was identified as high-risk if >=1 mutation in the stemness feature was detected in their ctDNA. Finally, 16 EMT-related mutated genes were identified: *FGFR1*, *FGFR2*, *RPS6KB1*, *IGF1R*, *MDM2*, *MAP2K4*, *ACIN1*, *BRIP1*, *EZH2*, *PPM1D*, *POLE*, *EXT1*, *EGFR*, *FAT1*, *MET*, *TJP1*. The presence of these mutations in ctDNA indicated shorter progression-free survival (PFS) and higher recurrence rates.

### Trajectory analysis

R package *monocle3* (version 0.1.2) was used to infer cell differentiation trajectory[48]. Pesudotime analysis was performed based on the learned trajectory to explore impressive translational cell relationships. This package comprehensively considered various requirements, including trajectory discontinuity, topology structure, and bifurcated trajectory, to select the most suitable trajectory-construction algorithm by function guidelines_shiny.

### Vascular proximity analysis

In the scSpatial-seq of all patients, the vasculature was annotated by professional pathologists based on HE-stained sections. In the vascular proximity analysis, the distance from each cell to the center of the nearest blood vessel was first calculated and identified as the vascular distance. Subsequently, the entire slice was divided into 50×50 spots. For each spot, the number of cells of various types, the average vascular distance, and the transcriptomic scores were calculated. Scores that showed a significant negative correlation with the vascular distance were considered to exhibit vascular proximity (200 genes, Pearson *P*.adj<0.05, *r* > 0.15).

### External validation based on H&E staining of breast cancer patients

To verify the association between vasculature and immune infiltration, we included 200 HE-stained images from patients covering all breast cancer subtypes. The images were segmented into patches of 250×250 μm. A random 25% of patches were selected from each slide, resulting in a total of 124,667 patches. After the segmentation was completed, using the nuclei segmentation dataset PanNuke[49], we annotated cell types into three categories: connectivity, inflammatory, and neoplastic. Specifically, the connectivity category represents the stromal area rich in vasculature, the inflammatory category represents the immune area with active immune responses, and the neoplastic category represents the epithelial area predominantly composed of malignant cells.

### Spatial pseudotime gradient

The single-cell spatial image was divided into a 20×20 grid of uniform squares, excluding areas devoid of cells, and the mean pseudotime for cells within each square was calculated. For each grid intersection, consider the four adjacent squares, and two pairs of diagonal squares form two perpendicular vectors. The resultant vector served as the spatial pseudotime gradient.

### Investigation and validation of B-cell-related epithelial mutations

To further investigate the epithelial genetic features associated with B immunity, we constructed a pipeline to explore and validate B-related mutations. Specifically, we first identified epithelial cells that showed significant interaction with B cells in scSpatial-seq data and obtained their transcriptomic characteristics. These characteristics were then transferred to scFAST-seq data at the same proportion using the AddModuleScore function. Subsequently, we identified differential mutations in these epithelial cells based on Fisher’s exact test. For each non-synonymous mutation, we transferred it back to scSpatial-seq data using the pipeline mentioned in the “Mutational label transfer on single gene from scFAST-seq to scSpatial-seq” section to explore the correlation between mutational scores and the number of B cells surrounding each epithelial cell. The validated mutations were then further investigated in the TCGA and METABRIC datasets to determine whether patients with these gene mutations exhibited higher B immunity.

### Identification of TLS

The identification of TLS is based on the spatial aggregation of T and B cells. Firstly, the numbers of T lineage and B lineage cells surrounding each cell were counted, and the product of these two numbers was used as the T-B-aggregation score. Subsequently, a density plot of the T-B-aggregation scores of immune cells was constructed. The threshold was set at two standard deviations to the right of the peak, and cells with scores above this threshold were considered to be part of TLS. Subsequently, we validated the characteristics of TLS from perspectives including pre-defined TLS score, cell proportion, immune function, and survival. In the analysis of epithelial cells, a density plot of the T-B-aggregation scores of epithelial cells was created. The threshold was also set at two standard deviations to the right of the peak, and cells with scores above this threshold were considered to be TLS-related epithelium.

### TME neighborhood analysis and patient stratification

Cells within a 250-micrometer peripheral range were assessed for the intensity of key interactions. Spatial interaction analysis from CellChat[50] was employed, incorporating all variable interaction pairs into the analysis. The TME neighborhood was delineated by identifying the types and quantities of non-epithelial cells around each epithelial cell. Hierarchical clustering of epithelial cells was performed using the hclust function. The determination of the cluster number was guided by the gap statistic. Five neighborhood-based clusters were subjected to DEG to obtain five transcriptomic scores, with the thresholds of *P*.adj <0.05 and log2 (fold change)>1, and an upper limit of 200 genes per score. Subsequently, in the patient stratification analysis based on METABRIC and TCGA, transcriptomic scores were transferred to the clinical cohort via the *GSVA* package (version 1.50.0)[51]. The group classification of each patient was ascertained by the highest of the five neighborhood scores. For survival analysis, the *survival* package was employed to generate Kaplan-Meier curves. Additionally, differential mutation analysis was carried out using Fisher’s exact test. All transcriptomic scores involved in this part were summarized in Supplementary Table 5.

### Survival analysis

The analysis was conducted using the *survminer* (version 0.4.9) and *survival* (version 1.5-7) packages, with a *p*-value threshold of less than 0.05 considered to indicate significance. Only data from estrogen receptor-positive (ER-pos) breast cancer cases were selected from the METABRIC dataset, whereas all breast cancer samples were included in the TCGA dataset.

### Statistics and reproducibility

Statistical analysis was conducted in R (v.4.3.3). Multiple testing correction was performed using the Benjamini–Hochberg (BH) method to generate all adjusted P-values (*P*.adj). For Pearson correlation, T-test, and Wilcoxon test analyses, the significance threshold was defined as *P*.adj less than 0.05. Numerical results were reported as means ± SEM as indicated. The Mann-Whitney test assessed Statistical significance between means (GraphPad Prism 9; GraphPad Software Inc., La Jolla, CA).

## Results

### Single-Cell Multi-Omics Data Integration in Luminal Breast Cancer

We focused on the luminal subtype of breast cancer, one of the most common cancer types[1, 52, 53], and collected matched tumor tissues and peripheral blood samples (Fig. 1A). To ensure sufficient ctDNA and exclude confounding effects of distant metastatic niches, we enrolled ten patients with non-metastatic high-risk hormone receptor-positive/human epidermal growth factor receptor 2-negative (HR^+^/HER2^-^) breast tumors characterized by features associated with elevated ctDNA release, including a mean tumor size >3 cm[54], lymphovascular invasion[55], and Ki67 ≥20%[56] (Fig. 1B; Table S1). Tumor tissues were analyzed for single-cell mutations and transcriptomes (scFAST-seq)[22, 23], single-cell spatial transcriptome (scSpatial-seq)[24], and bulk genomic alterations (tDNA sequencing). From blood samples, two DNA components were extracted: the germline genomic profile from peripheral blood mononuclear cells (PBMCs) and the mutation profiles of cell-free DNA (cfDNA), both sequenced using the same gene panel as the tumor DNA to ensure cross-modality comparability (641 genes, 2.126 Mb; Data S1). The mean sequencing depths were 5,067× for tDNA and 6,451× for cfDNA, enabling mutation detection at a variant allele frequency (VAF) of 0.1%. Furthermore, all validation datasets enrolled in this study are summarized in Table S2.

**Fig. 1.**
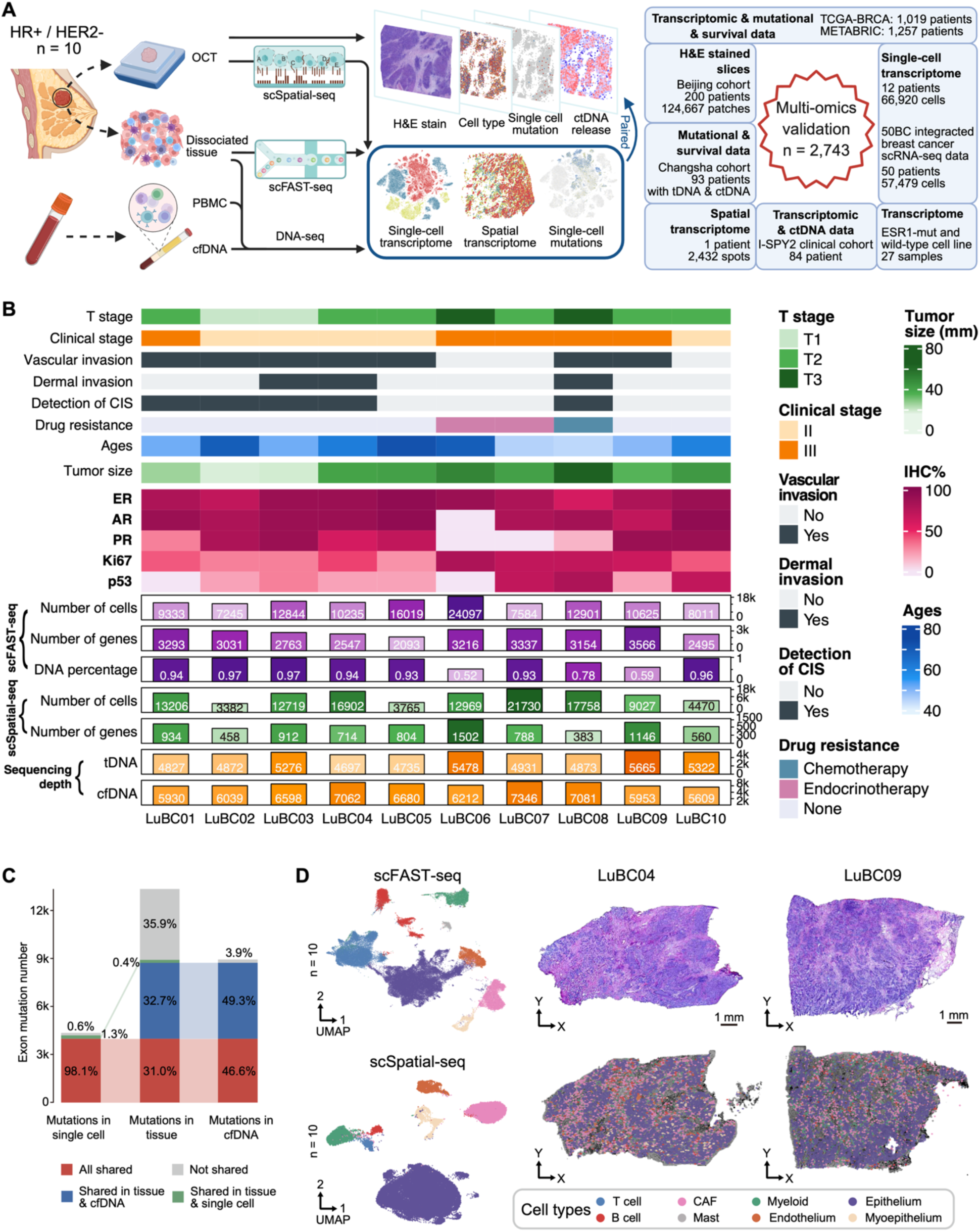
Study design and integration of single-cell multi-omics data. **(A)** Schematic of tissue sampling, sequencing workflow, and multi-omics validation. **(B)** Clinical profiles and sequencing metrics for the LuBC cohort (n = 10). **(C)** Comparison of exonic mutations across single-cell, bulk tissue, and cfDNA samples using a 641-gene panel (Supplementary Data 1). **(D)** UMAP visualization of cell subtypes (left) and spatial mapping with matched H&E staining for two representative samples (right). Abbreviations: HR+, Hormone Receptor-positive; OCT, optimal cutting temperature compound; PBMC, peripheral blood mononuclear cell; scSpatial-seq, single-cell spatial sequencing; scFAST-seq, full-length RNA sequence transcriptome sequencing.

The scFAST-seq was employed to obtain full-length single-cell transcriptomes and to infer single-cell mutations through reverse transcription (Fig. S1A)[22, 23]. To validate the reliability of single-cell mutations, we compared exonic mutations detected from the scFAST-seq with tDNA. Notably, 98.8% of the mutations in single-cell transcriptomes were confirmed in tDNA (Fig. 1C), with somatic mutation (PBMC < 1%) frequencies highly concordant between scFAST-seq and tDNA (Fig. S1F, Pearson *r* = 0.72, *P* < 0.001), with no detectable association with single-cell gene expression levels (*P* = 0.34). Furthermore, 97.5% of these single-cell mutations coexisted in both tissue DNA and cfDNA, confirming the accuracy of mutation detection in scFAST-seq (Fig. 1C). For spatial analysis, we used slide-labeled scSpatial-seq, in which single-cell nuclei were labeled with 1.5-µm barcode units. The transcriptomic data were then captured using a droplet-based scRNA-seq system (Fig. S1B)[24]. All quality control (QC) metrics and corresponding plots were provided in Data S2.

After QC and integration of scFAST-seq data from all patients, we retained 118,894 single-cell transcriptomes, which were categorized into eight major subgroups (Fig. 1D and Fig. S1D). Despite heterogeneous cell type distributions, all patients encompassed all major cell types (Fig. S1E). Utilizing the *inferCNV*[46], epithelial cells had abundant copy number variations (CNVs) across patients, showing high malignancy and inter-patient heterogeneity (Fig. S1G).

For multi-omics integration, we employed a transformer-based deep neural network to transfer cell type annotations from scFAST-seq to scSpatial-seq[21]. To validate transfer performance, we selected non-malignant cells with well-established markers and compared the transferred labels with manual annotations. The confusion matrix showed >90% of cells on the diagonal, with F1 scores >0.65 for all cell types. UMAP visualization revealed highly similar spatial distributions between the two annotations (Fig. S2A-C). These cell type annotations were subsequently transferred to the scSpatial-seq data, revealing distinct spatial distribution patterns for each cell type (Fig. 1D).

### Functional Diversity and Spatial Distributions of Malignant Epithelial States

We utilized a meta-cluster approach to uncover shared functional states among malignant cells and identified eight recurrent states (Fig. 2A and 2B). The antigen presentation (AP) state showed elevated MHC molecules (*CD74*, *HLA*) linked to enhanced antigen presentation. The mesenchymal progenitor-like (Mes) state, marked by *VIM*, *EGFR*, and *PTN*, exhibited activated epithelial-mesenchymal transition (EMT) and stemness pathways. The Mitotic state featured high *MKI67* expression with active mitotic checkpoint signaling. The Stress state highly expressed apoptosis-and chaperone-related genes (*EGR1*, *FOS*, *MYC*). The remaining states included ECM (*COL*/*WNT*), oxidative phosphorylation (OP, mitochondrial activity), mammary gland (Gland), and estrogen response (ERresp, high *ESR1* expression).

**Fig. 2.**
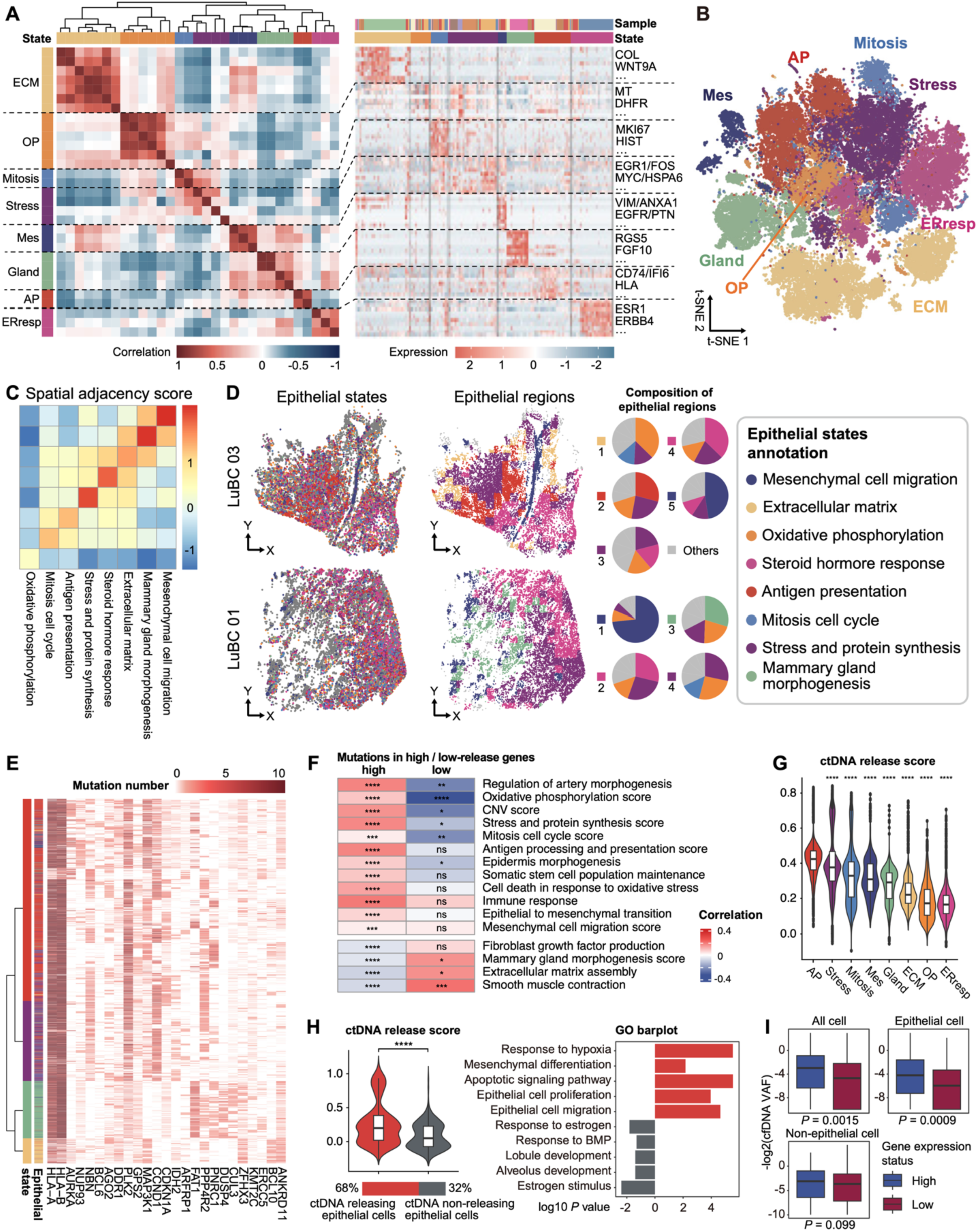
Malignant epithelial states and factors associated with ctDNA release. **(A)** Heatmap of eight epithelial states identified by gene signatures. **(B)** t-SNE projection of states. **(C)** Spatial adjacency scores between identified states. **(D)** Spatial mapping of epithelial states and corresponding regions for two representative samples (left), with pie charts indicating state compositions (right). **(E)** Differential functional mutation counts across epithelial states. **(F)** Correlations between biological pathways and somatic mutations within ctDNA-high and ctDNA-low release gene sets. **(G)** ctDNA release scores across epithelial states. **(H)** Comparison of ctDNA release scores (left) and enriched GO terms (right) between ctDNA-releasing and non-releasing epithelial cells. **(I)** Distribution of cfDNA VAF across cell types, stratified by gene expression levels in Changsha cohort. Abbreviations: ECM, extracellular matrix; OP, oxidative phosphorylation; Mes, mesenchymal cell migration; Gland, mammary gland morphogenesis; AP, antigen presentation; ERresp, steroid hormone response; VAF, variant allele frequency. Statistics: Adjusted P values from Wilcoxon (G) or two-tailed Student’s t test (I); ns, not significant; * *P* < 0.05; ** *P* < 0.01; *** *P* < 0.001; **** *P* < 0.0001.

Although all states were contributed by multiple patients, several states (AP, Gland, ECM, ERresp) showed high intra-patient heterogeneity (Fig. S2D). Nevertheless, core markers and functional pathways remained consistent between cells from dominant and other patients (Fig. S2E and S2F). Application of this eight-state framework to an external validation cohort of 50 breast cancer patients[57, 58] successfully classified 93% of cells, with all states represented by multiple patients and showing conserved marker expression and functional pathways, confirming external reproducibility (Fig. S2G-S2K).

Epithelial state labels from scFAST-seq were subsequently transferred to the scSpatial-seq data. To ensure reliable label transfer, we validated the biological identities of all states after integration. State scores identified from scFAST exhibited specific enrichment and ranked first in their corresponding states in scSpatial data (Fig. S3D). To quantify transfer accuracy, we performed a permutation test with 1,000 random label shuffles. For each state, the true transferred score was significantly higher than all permuted scores (*P.adj* < 0.001). Besides, key markers and functional pathway scores maintained similar patterns between the two platforms (Fig. S3A-S3C), confirming that transcriptional programs and functional states were faithfully transferred.

We then quantified spatial adjacency between epithelial states using average intercellular distances across all patients (Fig. 2C). Cellular contributions of individual patients are explicitly shown in Fig. S3E. Aggregate analysis of the full cohort revealed consistent core spatial associations. The Mes state clustered closely with Gland, ERresp, and ECM states in most patients (7/10: LuBC01,03,05,06,07,09,10), ERresp states were adjacent to Stress states (6/10: LuBC01,03,05,06,08,09), while the AP, Mitotic, and OP states exhibited weak spatial correlations with other states. Unsupervised clustering based on spot-wise state composition further identified specific spatial distributions (Fig. 2D). For instance, LuBC03 displayed a longitudinal intraductal Mes dominance, with a right region showing ERresp dominance while the left region exhibited mixed ERresp/Stress/AP/OP patterns. LuBC01 and LuBC08 both showed similar patterning, with discrete Mes-, ERresp-, and Stress/Mitosis-dominant regions (Fig. 2D and Fig. S3F). For the remaining seven patients, spatial patterning was more heterogeneous, yet all exhibited similar patterns to the cohort-wide trends (Supplementary Data 3). Collectively, these findings reveal spatial relationships between malignant states in luminal breast cancer, with patient-specific heterogeneity.

### Featured Mutational Profiles of Different Malignant Cell States

We first calculated tumor mutation burden (tTMB) and blood TMB (bTMB) based only on non-synonymous mutations, following thresholds below: PBMC VAF < 1% to eliminate germline contamination, cfDNA VAF ≥0.1%, and tDNA VAF ≥5%. We observed a median tTMB of 6.32 Muts/Mb, a 62.8% mutation retention rate in cfDNA, and a median bTMB of 3.06 mut/Mb (Table S2). This bTMB falls within the normal range reported in previous studies using targeted panel sequencing[59], ensuring the biological plausibility of our mutation data.

To investigate the effects of somatic mutations on epithelial cell functions, we analyzed the single-cell mutation profiles in the scFAST-seq data across all patients. Mutations were required to be present in tDNA and absent in PBMC to ensure their somatic origin. Because mutations were present in tDNA, which is not subject to RNA editing, this strategy inherently avoids false positives derived from RNA editing. First, we compared differential mutation numbers among specific cell populations and identified functionally different mutations in each epithelial state (Fig. 2E). Different cell populations exhibited distinct mutational genes. Compared to other epithelial states, the AP state had the highest number of mutations and mutated cells (Fig. S3G). Furthermore, a significant positive correlation was found between antigen presentation pathways and overall mutation burden in cells of AP state (Fig. S3H).

### Single-Cell ctDNA Tracing and Key Factors Associated With ctDNA Release

To obtain ctDNA release profiles for single cells, we compared mutation frequencies between tDNA and ctDNA. Accounting for patient-specific ctDNA dilution, we calculated the mean dilution rate across all mutations (cfDNA VAF/tDNA VAF; Fig. S4A) for each patient and then identified genes with cfDNA enrichment significantly exceeding this background (t-test, *P*<0.05). Notably, non-functional mutations, including synonymous and non-coding mutations, were retained in the analysis to identify sufficient biomarkers for ctDNA tracing.

At the gene level, we identified 17 genes with significantly higher mutation frequencies in ctDNA (ctDNA-high) and 20 genes with lower mutation frequencies (ctDNA-low; Table S4). ctDNA-high genes were primarily involved in cell growth, apoptosis, and damage repair, while ctDNA-low genes were enriched in immune regulatory pathways (Fig. S4B). Notably, VAFs of biomarker mutations in ctDNA and tDNA showed no evident patient bias (Fig. S4C). At the single-cell level, we calculated the somatic mutation rates in the ctDNA-high and ctDNA-low gene panels to evaluate cellular ctDNA contribution. Increased ctDNA release was associated with antigen presentation, cellular stress, and apoptosis, while negatively correlated with the ECM pathway and normal mammary function (Pearson |*r*| > 0.1, *P* < 0.05) (Fig. 2F).

By selecting genes positively correlated with ctDNA-high mutations, we derived a 77-gene signature and defined a ctDNA release score (Table S5). This score was validated through three independent approaches.

First, using scFAST-seq data, we validated the score using mutations not included in the biomarker selection. Cells with ctDNA-detectable mutations exhibited significantly higher ctDNA release scores than non-releasing cells (Wilcoxon test, *P* < 0.001; Fig. 2G and Fig. S4D). Only 68% of malignant cells harbored ctDNA-detectable mutations. These releasing cells exhibited enhanced proliferation, stemness, and hypoxia pathways with reduced normal mammary function (Fig. 2G). Second, external validation in the pre-treatment data of the I-SPY2 breast cancer cohort[38] (n = 84) confirmed clinical relevance. ctDNA-positive patients showed higher ctDNA release scores than ctDNA-negative patients (0.315 vs. 0.189, Wilcoxon test, *P* < 0.05; Fig. S4E). Moreover, correlations between ctDNA release and key biological factors, including proliferation, stress, and apoptosis, were fully recapitulated (Fig. S4E). Third, to ensure robustness against patient-specific bias, we performed leave-one-out validation. Sequential exclusion of each patient yielded stable correlations between the modified ctDNA release score and both survival (hazard ratio variation < 15%; all log-rank *P* < 0.05) and biological features (Fig. S4F), confirming intra-patient robustness.

We next used ctDNA release score to compare ctDNA release for different states. The Stress, Mes, AP, and Mitosis states exhibited high ctDNA release levels, while ECM, ERresp, OP, and Gland states showed lower levels (Fig. 2H). These findings were validated using several datasets, including scSpatial-seq, Jia-2023[25], and TCGA (Fig. S4G).

We further examined the association between gene expression and ctDNA release for each gene. In epithelial cells, genes with higher expression were associated with higher mutation read counts in ctDNA (Fig. S4I). Besides, cells were classified into epithelial, immune, and stromal to identify highly expressed genes within each population.

We then compared their ctDNA mutation frequencies in the Changsha cohort. Only genes highly expressed in epithelial cells showed significantly elevated ctDNA mutation rates (Fig. 2I and Fig. S4H). These results indicated that the abundance of a gene in ctDNA is positively associated with its expression level in malignant epithelial cells[60].

### ctDNA Non-Releasing *ESR1* Mutation Was Linked to Enhanced Mammary Functions and Reduced Immune Infiltrations

The above findings linked ctDNA release to malignant functional state. However, whether mutation detectability in ctDNA depends on cellular context remains unclear. We therefore examined *ESR1*, a canonical driver of HR^+^ breast cancer, to assess the relationship between *ESR1* mutations and ctDNA release. Of note, there was only one somatic mutation in the *ESR1* gene among ten samples, consistent with its overall mutation rate[61]. In LuBC07, the *ESR1* mutation rates (NM_000125.4: c.A1613G; p. D538G) were 30.1% in tissue, 12.8% in scFAST-seq data, but undetectable in ctDNA (Fig. 3A). Mutated tumor cells displayed lowered *ESR1* expression but elevated ESR1-related pathways, consistent with the constitutive activation of ESR1 pathways caused by *ESR1* mutations [62] (Fig. 3B and Fig. S5A).

**Fig. 3.**
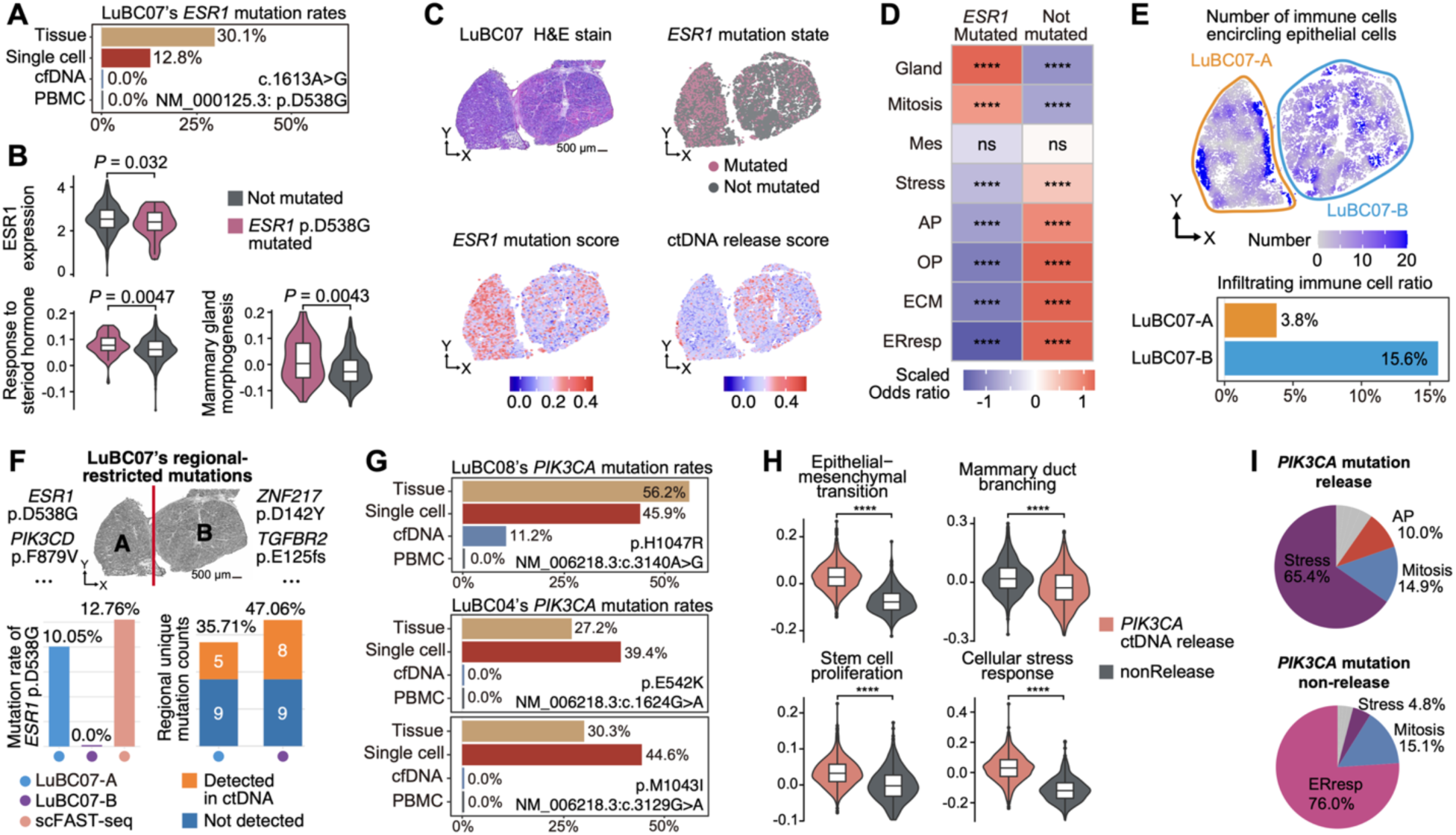
Multi-omic integration reveals spatial and functional heterogeneity of *ESR1* and *PIK3CA* mutation-associated ctDNA release. **(A)** Mutation rates of *ESR1* (NM_000125.3:p.D538G) across tissue, single-cell, cfDNA, and PBMC samples in LuBC07. (B) *ESR1* expression, hormone response, and mammary morphogenesis across different *ESR1* mutation statuses. **(C)** Spatial visualization of *ESR1* mutation status, *ESR1* mutation score, and ctDNA release score in LuBC07. **(D and E)** Distribution of (D) epithelial states and (E) immune cells in LuBC07. **(F)** Region-restricted mutations in LuBC07 (top); mutation rates of *ESR1* p.D538G across LuBC07-A, LuBC07-B, and scFAST-seq (left); ctDNA-detection rates of region-restricted mutations (right). (G) *PIK3CA* mutation rates at three loci across bulk tissue, single-cell, cfDNA, and PBMC samples in LuBC04/08. **(H and I)** (H) Pathway scores and (I) epithelial state compositions of *PIK3CA*-releasing and non-releasing epithelial cells. Statistics: Adjusted *P* values from two-tailed Student’s t-test; ns, not significant; **** *P* < 0.0001.

To investigate the spatial distribution of *ESR1* D538G, we developed an *ESR1* mutation score (200 genes; Supplementary Data 4) to map *ESR1* mutational status in scSpatial, preserving the 12.8% frequency in scFAST-seq. Mutated cells were significantly enriched in the left region of LuBC07, indicating divergent evolutions between regions (Fig. 3C). Trajectory analysis confirmed this finding, with *ESR1* mutations enriched in the left branch (Fig. S5B and S5C). The *ESR1* mutation-enriched region was dominated by the ctDNA-low-release Gland state (Fig. 3D), exhibiting low apoptosis, low stress, and low immune infiltration, likely shielded by an immunoregulatory rim (Fig. 3E and Fig. S5D and S5E). Spatial regional sequencing validated the heterogeneous distribution of *ESR1* D538G (left VAF = 10.05%, absent in right), and further revealed a lower ctDNA detectability in the left region (35.71% vs 47.06%) (Fig. 3F).

We further performed external validation using bulk RNA-seq data from cell lines harboring *ESR1* D538G[41]. Cell lines carrying *ESR1* D538G exhibited significantly higher *ESR1*-mut score (Wilcoxon test, *P* = 0.024) but lower ctDNA release scores (Wilcoxon test, *P* = 0.031), accompanied by enhanced mammary functions and decreased apoptosis pathways (Fig. S5F and S5G). This validation in a purely epithelial culture directly demonstrated that *ESR1* D538G may intrinsically reduce ctDNA release through mutation-associated function alterations[63]. Our findings establish associations between cell states, spatial contexts, and ctDNA release kinetics in one patient. These correlations generate strong hypotheses for future mechanistic investigation.

### *PIK3CA* Mutations Showed Divergent Associations With ctDNA Release

*PIK3CA* mutations are among the most frequent somatic alterations in breast cancer[64]. We identified three *PIK3CA* mutations with divergent ctDNA detectability: the NM_006218.3:c.3140A>G:p.H1047R kinase domain mutation was detected in ctDNA[65], whereas the NM_006218.3:c.1624G>A:p.E542K helical domain mutation and the NM_006218.3:c.3129G>A:p.M1043I C-terminal mutation[66] were not (Fig. 3G).

Mutant cells with ctDNA-releasing mutations demonstrated more pronounced malignant characteristics, including EMT, stemness, and cellular stress (Fig. 3H, Wilcoxon test, *P* < 0.001). This result is supported by experimental evidence showing that H1047R has higher tumor-promoting ability[65]. Notably, the dominant epithelial state in cells with releasing *PIK3CA* mutations was Stress (a ctDNA-high-release state), whereas for cells with non-releasing mutations was ERresp (a ctDNA-low-release state) (Fig. 3I). This contrast mirrored the pattern of *ESR1* D538G in LuBC07: while *ESR1* D538G drove a low-release Gland state, ctDNA-releasing *PIK3CA* mutations promoted stress and stemness. Together, these findings reinforced that mutation-associated functional states play a critical role in shaping ctDNA release heterogeneity. Different *PIK3CA* mutations showed divergent associations with ctDNA release and functional states, but validation in larger cohorts is required to draw definitive conclusions.

### Mutation Spatial Heterogeneity Associated With ctDNA Release

Beyond mutation-intrinsic effects, we examined whether the spatial distribution of mutations contributes to ctDNA release heterogeneity. Spatial regional sequencing (Supplementary Data 5) revealed that a substantial proportion of somatic mutations (20.7%–47.0% per patient) were detected exclusively in a single region, exhibiting spatial heterogeneity (Fig. S5H). ctDNA detectability varied markedly across regions, with significant differences even within the same patient (chi-square test, *P* < 0.05; Fig. S5I). Importantly, region-restricted mutations showed significantly lower ctDNA detectability than commonly distributed mutations (*P* = 0.002), and the proportion of region-restricted mutations negatively correlated with ctDNA detectability across regions (Pearson r = -0.43, *P* = 0.028) (Fig. S5J and S5K). These findings indicated the association between the spatial heterogeneity of mutations and ctDNA release.

### Spatial Regions Associated With EMT and ctDNA Characteristics

Next, we further examined how spatial distribution influences ctDNA release, focusing on the Mes state, characterized by EMT features and high ctDNA release potential. First, we examined its spatial distribution. Spatial adjacency analysis showed that Mes state, which serves as an early point in trajectory analysis, formed distinct spatial clusters rather than being dispersed among other states (Fig. S6A-S6C). We further defined spatial EMT regions as areas (radius 250 µm) with significant Mes-cell enrichment across the full cohort (Fig. 4A, 4B and Fig. S6D). The patient contribution to the subsequent analysis is quantified in Fig. S6E.

**Fig. 4.**
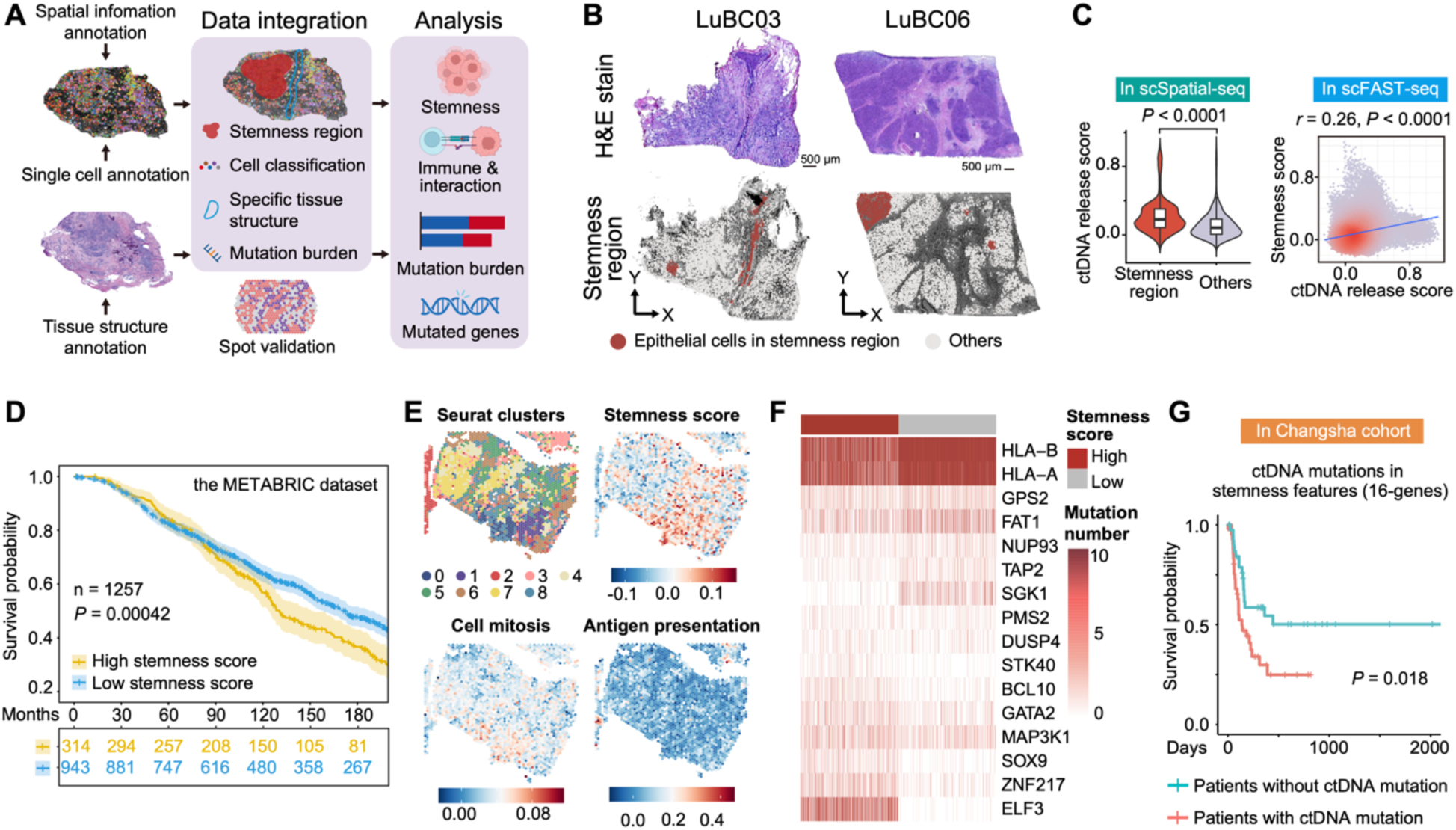
EMT-associated features modulates spatial ctDNA release and predicts poor clinical outcomes. **(A)** Overview of analytical frameworks for integrating single-cell multi-omics with spatial transcriptomics. **(B)** Spatial distribution of spatial EMT regions in two representative patients with matched H&E staining. **(C)** Comparison of ctDNA release scores between epithelial cells within vs. outside of regions in scSpatial-seq data (left), and correlation between spatial EMT scores and ctDNA release scores in scFAST-seq data (right; Pearson’s *r* = 0.26, *P* < 0.0001). **(D)** Kaplan-Meier survival analysis of patients in the METABRIC dataset, stratified by spatial EMT scores (*P* = 0.00042, log-rank test). **(E)** Spatial mapping of external ER+ breast cancer tissue sections, colored by cell clusters and corresponding molecular scores. **(F)** Mutation counts of differentially mutated genes in epithelial cells, grouped by high vs. low scores. **(G)** Progression-free survival (PFS) of the Changsha cohort, stratified by the presence or absence of ctDNA mutations in spatial EMT scores (*P* = 0.018, log-rank test).

Notably, significantly elevated ctDNA release scores were observed within spatial EMT regions (Fig. 4C). Epithelial cells within these regions exhibited reduced immune infiltrations, elevated hallmarks of hypoxia and proliferation, and enhanced stemness-related genes (Fig. S6F and S6G). Further differential expression analysis between cells inside and outside regions yielded a spatial EMT score (90 genes; Supplementary Data 4), predominantly expressed in epithelial cells (Fig. S6H). Importantly, a significant positive correlation was observed between the score and the ctDNA release score in scFAST data (Fig. 4C). High spatial EMT scores predicted worse overall survival in the METABRIC cohort[28] (Fig. 4D). External validation in an independent spatial transcriptome dataset[26] confirmed that high-scoring spots were enriched for EMT, cell division, and stemness, whereas low-scoring spots were associated with immune response (Fig. 4E).

To investigate the ctDNA underpinnings of the spatial EMT region, we performed differential mutation analysis on scFAST cells with high spatial EMT scores, identified mutations in epithelial proliferation and DNA repair pathways (Wilcox test *P.adj* < 0.05, Fig. 4F and Fig. S6I). To further assess its clinical relevance, we investigated the correlation between ctDNA mutations in the spatial EMT score and breast cancer relapse in our Changsha cohort, identifying a 16-gene mutation panel. Notably, mutations in the 16-gene panel strongly predicted shorter progression-free survival (Fig. 4G; Table S6). This suggested that the mutational composition of ctDNA, particularly mutations linked to EMT and stemness, can provide prognostic information.

### Intraductal Luminal Progenitors Exhibited Enhanced ctDNA Release Potential

Beyond tumor-intrinsic spatial organization, spatial architectures within tumor also impact ctDNA release. The breast duct represents a key structural component in luminal breast cancer, yet its immune interactions and ctDNA release remain poorly understood[67]. Breast ductal architectures were identified by pathologists in three patients (LuBC01/03/04, Fig. S7A). Mes-state tumor cells were concentrated within breast ducts, enclosed by MYH11-positive myoepithelial layers[67] (Fig. 5A). This distinct architecture was consistent with ductal carcinoma *in situ* structure[68]. Importantly, these luminal cells exhibited strong stemness features and high expression of the progenitor marker KIT[67] (Fig. 5A). Trajectory analysis revealed that Mes state exhibited the earliest pseudotime and was closest to the breast duct, confirming its luminal progenitor identity (Fig. 5B).

**Fig. 5.**
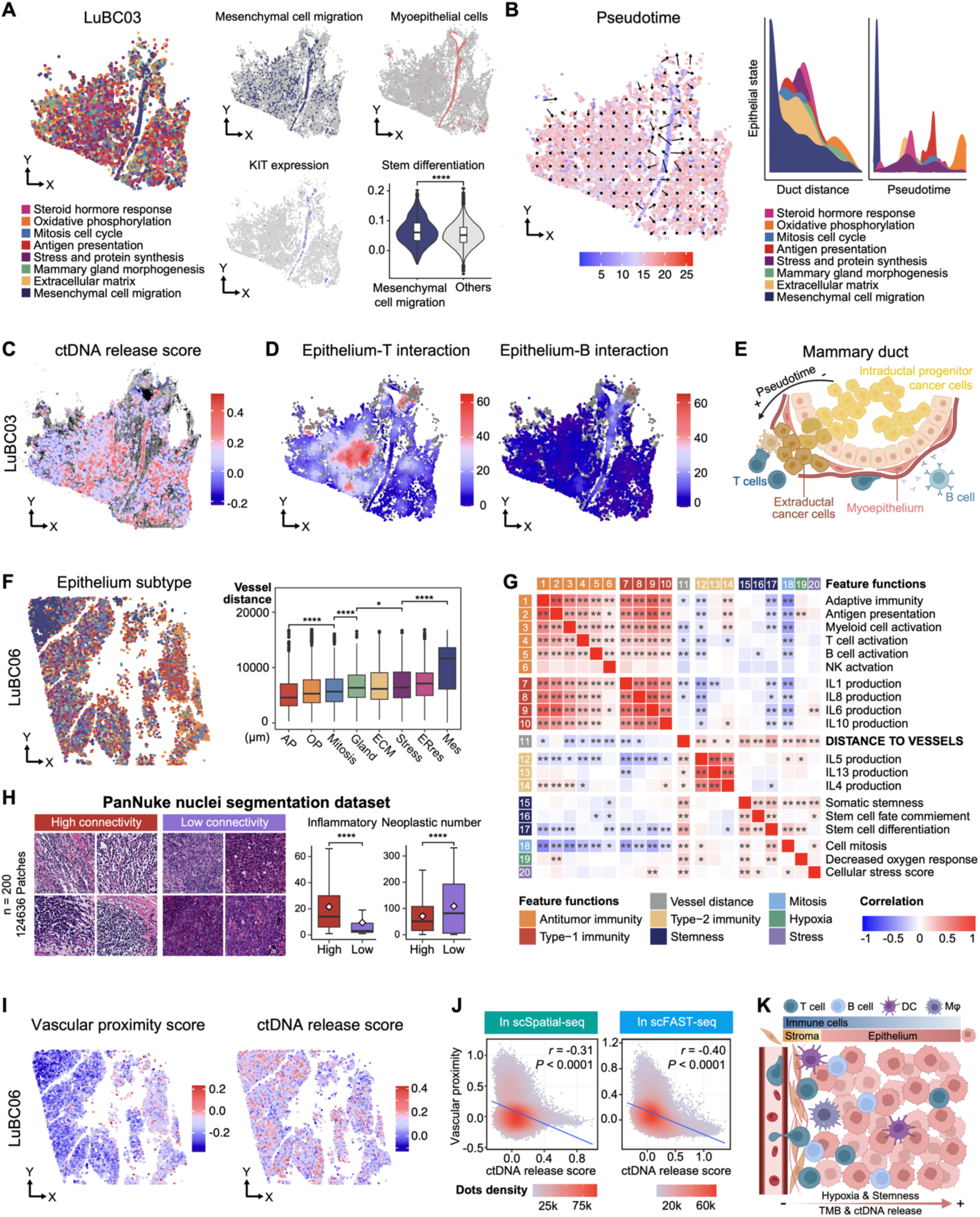
Ductal and vascular structures shape the tumor microenvironment and modulate ctDNA release. **(A)** Spatial visualization of epithelial states, myoepithelial cells, and KIT expression, with stem differentiation scores in the Mes state. **(B)** Spatial pseudotime gradient (left) and epithelial state dynamics along duct distance and pseudotime (right). **(C)** Spatial mapping of ctDNA release scores in a representative patient. **(D)** Spatial mapping of epithelial– immune cell interaction scores in a representative sample. **(E)** Schematic of the ductal carcinoma in situ to invasive carcinoma progression. **(F)** Spatial distribution of epithelial states (left) and distances from blood vessels (right). **(G)** Correlation between 19 feature functions and vascular distance. **(H)** H&E staining (left) and quantification of inflammatory scores and neoplastic cell counts (right) grouped by high or low connectivity patches (PanNuke nuclei segmentation dataset). **(I)** Spatial mapping of vessel and ctDNA release scores for a representative sample. **(J)** Negative correlation between vascular proximity and ctDNA release scores in scSpatial-seq (left; Pearson’s *r* = -0.31, *P* < 0.0001) and scFAST-seq (right; Pearson’s *r* = -0.40, *P* < 0.0001). **(K)** Schematic of the perivascular TME. Statistics: Adjusted *P* values from two-tailed Student’s t-test; * *P* < 0.05; ** *P* < 0.01; *** *P* < 0.001; **** *P* < 0.0001.

Projection of the ctDNA release score onto scSpatial data revealed that intraductal luminal progenitors exhibited high ctDNA release potential (Fig. 5C). This finding validated the inclusivity of ctDNA for early subclones in tumor evolution, highlighting the potential of ctDNA for breast cancer early detection[13]. The key associations between Mes, KIT, stemness, and ctDNA release score were observed in all patients with ductal architectures (Fig. S7A-S7C).

### Intraductal Luminal Progenitors Mediate ctDNA Release by B Cell Recruitment

To investigate the immune interactions of luminal progenitors and their association with ctDNA release, we conducted spatial immune interaction analysis (250 µm radius). Intraductal tumor cells displayed reduced epithelial-T but enhanced epithelial-B cell interactions (Fig. 5D and Fig. S7D), patterns also observed in LuBC01 (Fig. S7E). Luminal progenitors highly expressed CCL28, whose receptor CCR10 was enriched in plasma cells, consistent with the CCL28-CCR10 axis mediating B recruitment (Fig. S7F)[69]. Regarding antigen epitopes, differential mutation analysis identified mutations enriched in apoptosis, Wnt signaling, and EMT pathways, including *TGFBR2*, *ATRX*, and *TNFAIP3* (Fig. S7G). Spatial mapping showed four mutations had significantly higher B cell enrichment than the negative control (Fig. S7H), and TCGA data validated significant B cell enrichment in *ATRX*- or *TNFAIP3*-mutant tumors (Fig. S7I).

Interestingly, ctDNA release from luminal progenitors was associated with the functional state of recruited B cells. B cells proximal to malignant cells exhibited decreased normal B functions and increased TGF-β secretion (Fig. S7J). Notably, the Mes state showed high TGFBR2 expression and TGF-β receptor pathway activity, which positively correlated with ctDNA release score (Fig. S7K).

In contrast, tertiary lymphoid structures (TLS) identified in adjacent regions displayed classical immune effector functions, and epithelial cells near TLS showed reduced ctDNA release scores, suggesting that organized antitumor immune responses may modulate ctDNA release (Fig. S8A-S8E). In conclusion, luminal progenitors evade T-cell infiltration, interact with B cells with a TGF-β-secreting phenotype, and exhibit high ctDNA release potential (Fig. 5E).

### Vasculature Modulated TME and ctDNA Release Through Immune Infiltration and Hypoxia

In addition to the breast duct, blood vessels represent another key spatial architecture in luminal breast cancer. We next examined how vasculature shapes the TME and ctDNA release. Distinct vascular components were identified by pathologists in two patients, LuBC06 and LuBC08 (Fig. S9A). Notably, the AP and OP states localized near blood vessels, whereas the Stress and Mes subgroup cells were located far and exhibited higher hypoxia levels (Fig. 5F and Fig. S9B). For a more detailed analysis, we partitioned slices into 50×50 small patches and conducted correlation analysis between biological characteristics and vessel distances. Immune cell densities, anti-tumor immunity, and AP-state cells increased near blood vessels. Conversely, stemness and hypoxia-related stress decreased (Fig. 5G). These results were validated via an external cohort of 200 patients from the PanNuke nuclei segmentation dataset[49]. Patches with greater connectivity, indicating higher microvascular density, showed significantly more immune infiltration and fewer neoplastic cells (Fig. 5H).

We identified a gene set of 200 genes that positively correlated with vascular proximity as the vascular proximity score (Supplementary Data 4). This score showed a significant negative correlation with hypoxia and stemness, but a strong positive correlation with endothelial cell scores in invasive breast cancer[70] and bevacizumab response score[71] (Fig. S9C-S9E). Intriguingly, hypoxic regions distant from vessels in both patients exhibited higher ctDNA release (Fig. 5I and Fig. S9F). These findings were consistent in scFAST-seq and scSpatial-seq (Fig. 5J), further confirming the correlation between vascular distance and ctDNA release.

Transferring the vascular proximity score to scFAST-seq revealed that cells with low proximity scores carried more somatic mutations enriched in p53, DNA repairing, and UV-response pathways (Fig. S9G). In summary, proximity to vasculature defined a TME with strong immune infiltration and antigen presentation, whereas distant regions showed hypoxia, higher mutation burden, and increased ctDNA release (Fig. 5K). The higher ctDNA release in hypoxic regions distant from vessels, though counterintuitive, likely results from hypoxia-induced necrosis that releases DNA into circulation. Our results suggested that hypoxia-associated stress, not physical distance, is the key driver.

### Epithelial Cells Exhibited Distinct TME Interactions Based on Their Spatial Neighborhoods

Extending our analysis from specific spatial architectures to cellular neighborhoods, we performed single-cell spatial neighborhood analysis by identifying non-epithelial cells within a 250 μm radius of each epithelial cell[50]. Unsupervised clustering based on neighborhood composition classified epithelial cells into five clusters (Fig. 6A and 6B), with consistent immune infiltration patterns across all patients (Fig. 6C).

**Fig. 6.**
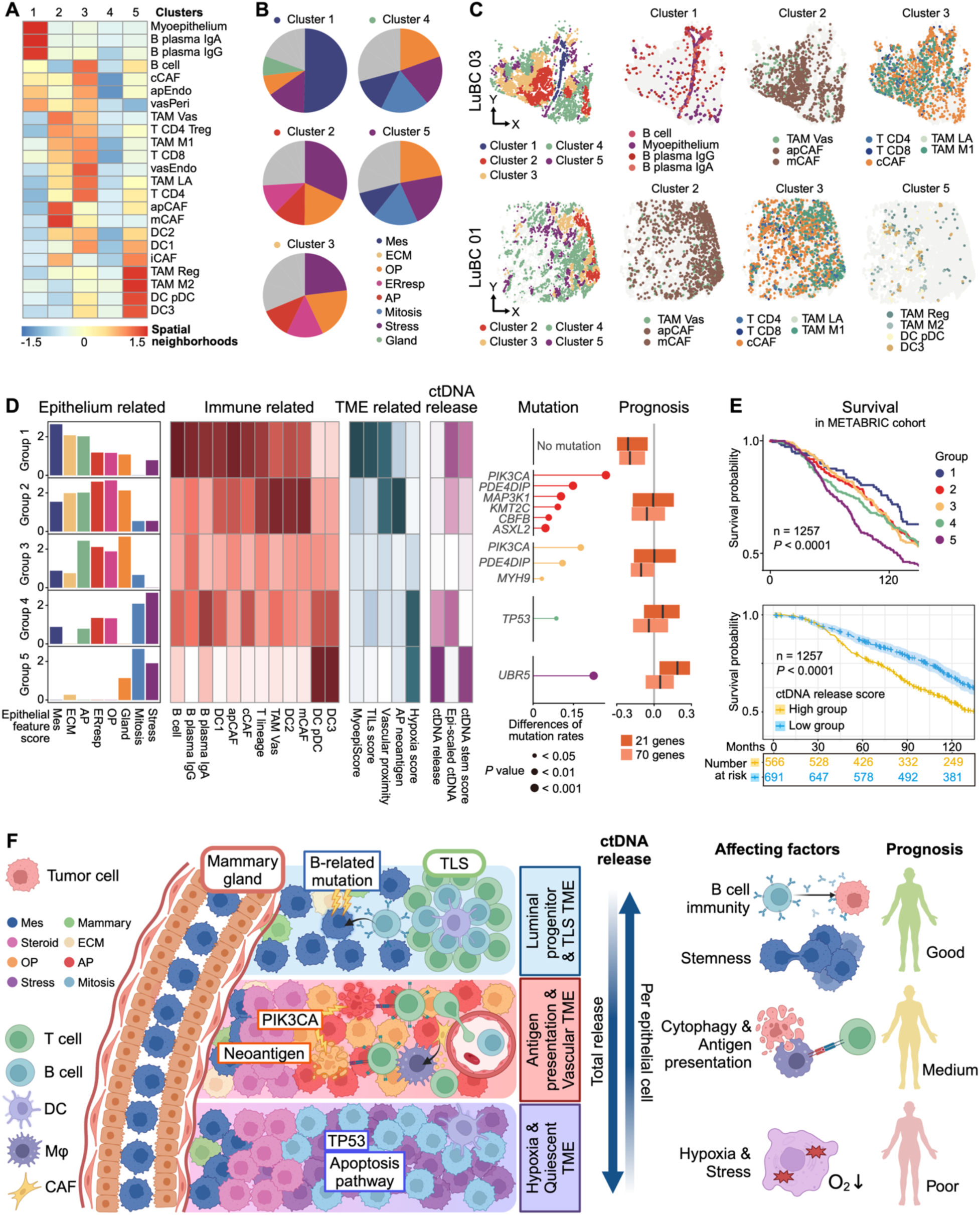
TME-based epithelial clustering and patient risk stratification for clinical prognosis. **(A)** Enrichment of non-epithelial spatial neighborhoods across five epithelial clusters. **(B)** Cellular compositions of epithelial states within each cluster. **(C)** Spatial mapping of epithelial clusters and representative immune cells. **(D)** Patient risk stratification in the METABRIC cohort (n = 1,904): (1) epithelial state features; (2–4) immune-related, TME-related, and ctDNA release scores; (5) mutation rates of differentially mutated genes; and (6) 21-gene and 70-gene scores. **(E)** K-M survival analysis in the METABRIC cohort for the five patient groups (top) and high vs. low ctDNA release groups (bottom). **(F)** Schematic of three distinct TMEs, their ctDNA release profiles, and prognostic implications. Abbreviations: TME, tumor microenvironment. Statistics: P values from log-rank test (E).

Among the five clusters, cluster 1, primarily composed of Mes state, was characterized by strong interactions with B cells and myoepithelium, along with prominent IL-10 and TGF-β receptor pathway activity (Fig. 6A and Fig. S10A). This cluster was present in four patients, including LuBC01/03/04 with breast ductal architectures (Fig. S10B). Clusters 2/3 showed similar profiles dominated by T cells, tumor-associated macrophages (TAMs), and cancer-associated fibroblasts (CAFs), with higher proportions of AP-state and enhanced immune recruitment. Cluster 5 was defined by immunomodulatory components, including M2 TAM (CD163, MRC1), regulatory TAM (CD40, CD274), DC3 (LAMP3, BIRC3), and pDC (TCF4, CLEC4C, CD274), while its epithelial composition resembled immune-quiescent cluster 4, both featuring Mitosis and Stress. Notably, cluster 4 accounts for the largest proportion of all epithelial cells. consistent with the limited immune infiltration characteristic of HR^+^ breast cancer (Fig. S10C).

Notably, clusters with the lowest anti-tumor immune infiltration (clusters 4 and 5) exhibited the highest ctDNA release scores (Wilcox test, *P.adj* < 0.001; Fig. S10D). Quantification across all epithelial cells further revealed a significant negative correlation between total immune neighbor count and ctDNA release score (Fig. S10E). However, not all immune cells exerted equivalent effects. Antitumor populations, including CD8^+^ T cells and M1 TAM, were associated with reduced ctDNA release. In contrast, immunomodulatory types, including regulatory TAM, M2 TAM, and DC3, were linked with increased ctDNA release scores (Fig. S10F). These findings aligned with our observations in ductal regions, while TGF-β-releasing B cells correlated with elevated ctDNA release (Fig. S7J and S7K), organized TLS with functional immune responses correlated to reduced release (Fig. S8C-S8E). In summary, the functional orientation of the immune microenvironment, rather than immune abundance alone, impacted ctDNA release.

### Clinical Utility of The TME Neighborhood Scores

To further investigate the clinical relevance, we derived five TME neighborhood scores (Supplementary Data 4) and applied them to METABRIC and TCGA cohorts via Gene Set Variation Analysis (GSVA)[51], classifying patients into five groups with consistent phenotypes across datasets (METABRIC in Fig. 6D; TCGA in Fig. S10G). Tumors in group 1 exhibited the highest Mes and ECM epithelial features, along with prominent B-cell infiltration and breast duct characteristics. Groups 2 and 3 showed strong AP/ERresp and immune enrichment, with group 2 displaying the most mutation and highest neoantigen presentation scores (genes correlating with mutation rates in AP state, 54 genes; Supplementary Data 4), and group 3 enriched for *PIK3CA* mutations and CAF/TAM accumulation (Fig. 6D). Groups 4 and 5 exhibited elevated Mitosis/Stress features, with group 4 being immune quiescent and group 5 being immunomodulatory (DC3/pDC).

For ctDNA release, highly malignant groups 4 and 5 demonstrated the highest ctDNA release scores. However, when adjusted for epithelial cells, the normalized ctDNA scores were highest in group 1, aligning with the ctDNA release potential of luminal progenitors. For prognosis, group 1 demonstrated significantly lower risk in both 21-gene[72] and 70-gene signature[73], while group 5 exhibited the highest risk (Fig. 6D). In the survival analysis, patients in group 5 had the poorest prognosis, followed by the group 4, groups 2 and 3, and group 1 with the best (Fig. 6E). Considering groups 4 and 5 also exhibited the highest overall ctDNA release, we then demonstrated that patients with higher ctDNA release signatures had significantly worse 5-year survival. This suggests that ctDNA can serve as a non-invasive biomarker to predict the prognosis of breast cancer patients, consistent with the recent findings from the monarchE study[74].

Notably, the patient risk stratification results recapitulated the findings from our TME neighborhood analysis. Patients with high antitumor immunity exhibited favorable survival outcomes but lower ctDNA release. In contrast, immune-quiescent or immunomodulatory patients displayed more malignant features, which were associated with elevated ctDNA release and poorer prognosis. These observations further indicated that ctDNA release reflects not simply tumor presence, but also the context of immune microenvironments[75].

## Discussion

Using HR⁺/HER2⁻ breast cancer as a model, we established an integrative analytical framework scMASTER that analyze ctDNA release at cellular level, dissected ctDNA release heterogeneity across transcriptional states, functional mutations, and spatial architectures, demonstrating that ctDNA release is shaped by both tumor-intrinsic programs and the functional orientation of TME.

Existing ctDNA origin-analyzing strategies have largely followed two paradigms, including methylation-based deconvolution which identifies tissue sources but cannot distinguish states within the same lineage[76], and clonal phylogenetic inference which requires serial sampling or ultra-deep WGS[77]. In contrast, scMASTER leverages co-detected somatic mutations as endogenous barcodes and directly links ctDNA release to specific malignant states. ctDNA-high mutations can be traced back to single-cell states to reveal the underlying biology; conversely, features identified from key cell states from scRNA-seq can guide the design of ctDNA panels. This framework requires standard inputs including tDNA sequencing, paired plasma cfDNA, and whole-length scRNA-seq. The pipeline is platform-agnostic and can be adapted to other gene panels or cancer types, offering a practical reference for broader liquid biopsy research.

Current understanding of ctDNA release has largely focused on production-side mechanisms such as apoptosis, necrosis, and active secretion[78]. As for the clearance side, ctDNA’s short half-life is primarily governed by Kupffer cell-mediated phagocytosis and circulating nucleases[79]. Recent work has shown that modulating in vivo clearance can alter ctDNA recovery by an order of magnitude[75], yet how production and clearance forces interact has remained unexplored at single-cell resolution. Our data directly evidence this net release balance. At scSpatial-seq, we observed that hypoxic zones distant from vasculature exhibited high ctDNA release, an apparent paradox that can be understood by the net release framework. On the production side, hypoxia-driven necrosis actively releases abundant DNA fragments[78]; on the clearance side, the sparse immune infiltration in these regions may reduce macrophage-mediated clearance[79]. Besides, elevated interstitial fluid pressure in avascular tumor regions may facilitate flow toward lymphatic vessels[80]. These observations indicate that ctDNA release is not merely a passive readout of tumor burden, but an integrated signal of both tumor biology and local immune surveillance.

The relationship between immunotherapy and ctDNA has drawn increasing attention[81]. Evidence suggested that the immune system plays a dual role by clearing ctDNA from circulation[79] or influencing its release through effects on tumor cells[82]. Recent clinical studies have shown that ctDNA dynamics can distinguish pseudoprogression from true progression during immunotherapy[83], underscoring that ctDNA levels reflect not only tumor shedding but also tumor immune status. Our spatial neighborhood analysis provided direct evidence for this bidirectional relationship. Epithelial cells surrounded by antitumor effectors exhibited lower ctDNA release scores, consistent with efficient immune-mediated tumor control or rapid clearance. In contrast, neighborhoods enriched for immunomodulatory components correlated with higher ctDNA release and enhanced stemness and stress features[84]. These findings may help explain the complex ctDNA dynamics observed during immunotherapy[85], as the absence of an expected ctDNA peak could reflect potent immune clearance. Collectively, these TME-centric findings reframe ctDNA dynamics as a real-time readout of tumor– immune interplay. This bidirectional lens provides a biological rationale for ctDNA-guided immunotherapy monitoring beyond tumor volume alone.

There are several limitations. To ensure the stability of cancer phenotypes, our analysis was restricted to HR^+^/HER2^−^ breast cancer. Expanding these findings to other subtypes or cancer types is crucial for understanding the heterogeneity of ctDNA release. Additionally, incorporating a broader range of multi-omics data, including methylation profiles, could facilitate a more comprehensive ctDNA tracing. Beyond the integration of one clinical cohort, multiple public datasets, and leave-one-out sensitivity analyses, further validation in larger, independent cohorts will be important to generalize these findings.

In conclusion, we constructed a single-cell multi-omics framework to trace ctDNA origins and systematically dissected release heterogeneity. By integrating transcriptomic, mutational, and spatial data, we revealed that ctDNA release is associated with different tumor-intrinsic functional states, modulated by functional mutations, and further shaped by spatial architectures as well as TME effects, ultimately linking ctDNA release heterogeneity to patient prognosis.

## Resource availability

### Lead contact

Requests for further information and resources should be directed to and will be fulfilled by the lead contact, Jiaqi Liu.

### Materials availability

This study did not generate new unique reagents.

### Data and Code Availability

The raw sequence data reported in this paper have been deposited in the Genome Sequence Archive in National Genomics Data Center, China National Center for Bioinformation/Beijing Institute of Genomics, Chinese Academy of Sciences (GSA-Human: HRA017586) that are publicly accessible at https://ngdc.cncb.ac.cn/gsa-human. The processed data are available under OMIX015902 in Open Archive for Miscellaneous Data at https://ngdc.cncb.ac.cn/omix. The code repository is publicly available at https://github.com/Pengming-Pu/scMASTER-single-cell-Mutational-and-Spatial-Transcriptomics-in-Enhanced-Resolution.

## Acknowledgments

We thank all the individuals, families, and physicians involved in the study for their participation.

## Author contributions

Conceptualization was performed by J.S., Y.Z., H.X., and J.L. Clinical biopsies were collected by B.L., Y.G., Z.J., C.G., and Y.L. Formal analysis was conducted by J.S., Y.Z., Y.H., and D.M. Experiments were carried out by P.P., C.G., B.L., and J.W. Bioinformatic analysis was performed by H.X., P.P., Y.Z., and J.L. Visualization was done by H.X., P.P., Y.Z., R.Z., and Z.G. The original draft was written by H.X., P.P., Y.Z., T.S., and J.L. Review and editing of the manuscript were completed by H.X., P.P., Y.Z., B.L., Y.G., H.C., Y.W., and J.L. Revisions were completed by H.X., P.P., C.G., Q.O., and J.W. Resources were provided by Q.O., J.L., H.X., B.L., and Y.W. Data curation was managed by H.X. and L.C. Project administration was overseen by J.L., X.W., and P.P.

## Ethics declarations

This study was reviewed and approved by the ethics committees at each participating hospital. Written informed consent was obtained from each participant.

## Supplemental information

**Fig. S1-S10**

**Table S1 |** Patient information and QC data.

**Table S2 |** The summary of all utilized datasets in this study.

**Table S3 |** The calculation of bTMB per patient.

**Table S4 |** The summary of biomarker selections.

**Table S5 |** ctDNA release score (77 genes).

**Table S6 |** The genes of relapse-associated mutated genes discovered in the Changsha cohort.

**Supplementary Data S1 |** Utilized targeted sequencing panel (641 genes, 2.126Mb).

**Supplementary Data S2 |** Key QC data for scFAST-seq and scSpatial-seq.

**Supplementary Data S3 |** Results of spatial proximity analysis of LuBC01-10.

**Supplementary Data S4|** All used transcriptomic scores in the manuscript.

**Supplementary Data S5 |** Spatial segmentation of FFPE tumor tissues from 10 patients for panel sequencing.

## References

1. Bray F, Laversanne M, Sung H, Ferlay J, Siegel RL, Soerjomataram I, Jemal A: Global cancer statistics 2022: GLOBOCAN estimates of incidence and mortality worldwide for 36 cancers in 185 countries. CA Cancer J Clin 2024, 74:229–263.

2. Mosele MF, Westphalen CB, Stenzinger A, Barlesi F, Bayle A, Bieche I: Recommendations for the use of next-generation sequencing (NGS) for patients with advanced cancer in 2024: a report from the ESMO Precision Medicine Working Group. Annals of Oncology 2024, 35:588–606.

3. Burstein HJ, Somerfield MR, Barton DL, Dorris A, Fallowfield LJ, Jain D: Endocrine Treatment and Targeted Therapy for Hormone Receptor-Positive, Human Epidermal Growth Factor Receptor 2-Negative Metastatic Breast Cancer: ASCO Guideline Update. Journal of Clinical Oncology 2021, 39:3959–3977.

4. Lloyd MR, Jhaveri K, Kalinsky K, Bardia A, Wander SA: Precision therapeutics and emerging strategies for HR-positive metastatic breast cancer. Nature Reviews Clinical Oncology 2024, 21:743–761.

5. Bhave MA, Quintanilha JCF, Tukachinsky H, Li G, Scott T, Ross JS: Comprehensive genomic profiling of ESR1, PIK3CA, AKT1, and PTEN in HR(+)HER2(-) metastatic breast cancer: prevalence along treatment course and predictive value for endocrine therapy resistance in real-world practice. Breast Cancer Research and Treatment 2024, 207:599–609.

6. Venetis K, Pepe F, Pescia C, Cursano G, Criscitiello C, Frascarelli C: ESR1 mutations in HR+/HER2-metastatic breast cancer: Enhancing the accuracy of ctDNA testing. Cancer Treatment Reviews 2023, 121:102642.

7. Zhang W, Brown EL, Usmani A, Earland N, Kang M, Olelewe C, Viswanathan A, Chauhan PS, Steen CB, Jeon HS, et al: Non-invasive profiling of the tumour microenvironment with spatial ecotypes. Nature 2026.

8. Gavish A, Tyler M, Greenwald AC, Hoefflin R, Simkin D, Tschernichovsky R, Galili Darnell N, Somech E, Barbolin C, Antman T, et al: Hallmarks of transcriptional intratumour heterogeneity across a thousand tumours. Nature 2023, 618:598–606.

9. Akcakanat A, Zheng X, Cruz Pico CX, Kim TB, Chen K, Korkut A: Genomic, Transcriptomic, and Proteomic Profiling of Metastatic Breast Cancer. Clinical Cancer Research 2021, 27:3243–3252.

10. Cohen SA, Liu MC, Aleshin A: Practical recommendations for using ctDNA in clinical decision making. Nature 2023, 619:259–268.

11. Nakamura Y, Ozaki H, Ueno M, Komatsu Y, Yuki S, Esaki T, Taniguchi H, Sunakawa Y, Yamaguchi K, Kato K, et al: Targeted therapy guided by circulating tumor DNA analysis in advanced gastrointestinal tumors. Nat Med 2024.

12. Pellini B, Chaudhuri AA: Circulating Tumor DNA Minimal Residual Disease Detection of Non-Small-Cell Lung Cancer Treated With Curative Intent. J Clin Oncol 2022, 40:567–575.

13. Panet F, Papakonstantinou A, Borrell M, Vivancos J, Vivancos A, Oliveira M: Use of ctDNA in early breast cancer: analytical validity and clinical potential. NPJ Breast Cancer 2024, 10:50.

14. Siravegna G, Lazzari L, Crisafulli G, Sartore-Bianchi A, Mussolin B, Cassingena A, Martino C, Lanman RB, Nagy RJ, Fairclough S, et al: Radiologic and Genomic Evolution of Individual Metastases during HER2 Blockade in Colorectal Cancer. Cancer Cell 2018, 34:148–162.e147.

15. Zeng Z, Yi Z, Xu B: The biological and technical challenges facing utilizing circulating tumor DNA in non-metastatic breast cancer patients. Cancer Letters 2025, 616:217574.

16. You Y, Fu Y, Li L, Zhang Z, Jia S, Lu S, Ren W, Liu Y, Xu Y, Liu X, et al: Systematic comparison of sequencing-based spatial transcriptomic methods. Nature Methods 2024.

17. Chung C, Yang X, Hevner RF, Kennedy K, Vong KI, Liu Y, Patel A, Nedunuri R, Barton ST, Noel G, et al: Cell-type-resolved mosaicism reveals clonal dynamics of the human forebrain. Nature 2024, 629:384–392.

18. Dang DK, Park BH: Circulating tumor DNA: current challenges for clinical utility. J Clin Invest 2022, 132.

19. Chen C, Guo Q, Liu Y, Hou Q, Liao M, Guo Y, Zang Y, Wang F, Liu H, Luan X, et al: Single-cell and spatial transcriptomics reveal POSTN(+) cancer-associated fibroblasts correlated with immune suppression and tumour progression in non-small cell lung cancer. Clin Transl Med 2023, 13:e1515.

20. Rodriguez-Meira A, Buck G, Clark SA, Povinelli BJ, Alcolea V, Louka E, McGowan S, Hamblin A, Sousos N, Barkas N, et al: Unravelling Intratumoral Heterogeneity through High-Sensitivity Single-Cell Mutational Analysis and Parallel RNA Sequencing. Mol Cell 2019, 73:1292–1305.e1298.

21. Yuan M, Wan H, Wang Z, Guo Q, Deng M: SPANN: annotating single-cell resolution spatial transcriptome data with scRNA-seq data. Brief Bioinform 2024, 25.

22. Sang G, Chen J, Zhao M, Shi H, Han J, Sun J, Guan Y, Ma X, Zhang G, Gong Y, et al: High throughput detection of variation in single-cell whole transcriptome through streamlined scFAST-seq. bioRxiv 2023:2023.2003.2019.533382.

23. Qiao N, Wang ZX, Zhang YL, Zhang LQ, Zhu HM, Weng XQ, Zhu YM, Cheng WY, Li JF, Jiang L, et al: HSCs/MPPs as cells of origin with altered differentiation hierarchy impairing immunomicroenvironment in PML::RARA and CBFα/β fusion AML. Proc Natl Acad Sci U S A 2026, 123:e2526334123.

24. Pan Y, Yan H, Han J, Wu R, Xu C, Lei G, Ma X, Guan Y, Li Z, Deng J, et al: Integrating single-nucleus barcoding with spatial transcriptomics via Stamp-seq to reveal immunotherapy response-enhancing functional modules in NSCLC. Cell Discov 2026, 12:10.

25. Jia Z, Xu H, Zhang Y, Cao H, Deng C, Xu L, Sun Y, Li J, Huang Y, Pu P, et al: Distinct discrepancy in breast cancer organoids recapitulation among molecular subtypes revealed by single-cell transcriptomes analysis. Clin Transl Med 2024, 14:e70023.

26. Wu SZ, Al-Eryani G, Roden DL, Junankar S, Harvey K, Andersson A, Thennavan A, Wang C, Torpy JR, Bartonicek N, et al: A single-cell and spatially resolved atlas of human breast cancers. Nat Genet 2021, 53:1334–1347.

27. Colaprico A, Silva TC, Olsen C, Garofano L, Cava C, Garolini D, Sabedot TS, Malta TM, Pagnotta SM, Castiglioni I, et al: TCGAbiolinks: an R/Bioconductor package for integrative analysis of TCGA data. Nucleic Acids Res 2016, 44:e71.

28. Curtis C, Shah SP, Chin SF, Turashvili G, Rueda OM, Dunning MJ, Speed D, Lynch AG, Samarajiwa S, Yuan Y, et al: The genomic and transcriptomic architecture of 2,000 breast tumours reveals novel subgroups. Nature 2012, 486:346–352.

29. Chen S, Zhou Y, Chen Y, Gu J: fastp: an ultra-fast all-in-one FASTQ preprocessor. Bioinformatics 2018, 34:i884–i890.

30. Dobin A, Davis CA, Schlesinger F, Drenkow J, Zaleski C, Jha S, Batut P, Chaisson M, Gingeras TR: STAR: ultrafast universal RNA-seq aligner. Bioinformatics 2013, 29:15–21.

31. Liao Y, Smyth GK, Shi W: The Subread aligner: fast, accurate and scalable read mapping by seed-and-vote. Nucleic Acids Res 2013, 41:e108.

32. Zheng GXY, Terry JM, Belgrader P, Ryvkin P, Bent ZW, Wilson R, Ziraldo SB, Wheeler TD, McDermott GP, Zhu J, et al: Massively parallel digital transcriptional profiling of single cells. Nature Communications 2017, 8:14049.

33. Lun ATL, Riesenfeld S, Andrews T, Dao TP, Gomes T, Marioni JC: EmptyDrops: distinguishing cells from empty droplets in droplet-based single-cell RNA sequencing data. Genome Biol 2019, 20:63.

34. Asmann YW, Parikh K, Bergsagel PL, Dong H, Adjei AA, Borad MJ, Mansfield AS: Inflation of tumor mutation burden by tumor-only sequencing in under-represented groups. npj Precision Oncology 2021, 5:22.

35. Parikh K, Huether R, White K, Hoskinson D, Mansfield AS: Tumor Mutational Burden From Tumor-Only Sequencing Compared With Germline Subtraction From Paired Tumor and Normal Specimens. JAMA Network Open 2020, 3:e200202.

36. Poh J, Ngeow KC, Pek M, Tan KH, Lim JS, Chen H, Ong CK, Lim JQ, Lim ST, Lim CM, et al: Analytical and clinical validation of an amplicon-based next generation sequencing assay for ultrasensitive detection of circulating tumor DNA. PLoS One 2022, 17:e0267389.

37. Yu Y, Jin M, Yuan W, Gong Y, Li S, Qin X, Hou J, Liu J, Liu S, Li H, et al: Engineered crRNA Drives RPA-T7-CRISPR/Cas14a Cascade for Ultrasensitive Detection of ctDNA PIK3CA H1047R. Adv Sci (Weinh) 2025, 12:e07126.

38. Magbanua MJM, Li W, Wolf DM, Yau C, Hirst GL, Swigart LB, Newitt DC, Gibbs J, Delson AL, Kalashnikova E, et al: Circulating tumor DNA and magnetic resonance imaging to predict neoadjuvant chemotherapy response and recurrence risk. npj Breast Cancer 2021, 7:32.

39. Butler A, Hoffman P, Smibert P, Papalexi E, Satija R: Integrating single-cell transcriptomic data across different conditions, technologies, and species. Nat Biotechnol 2018, 36:411–420.

40. Hao Y, Hao S, Andersen-Nissen E, Mauck WM, III, Zheng S, Butler A, Lee MJ, Wilk AJ, Darby C, Zager M, et al: Integrated analysis of multimodal single-cell data. Cell 2021, 184:3573–3587.e3529.

41. Yu L, Wang L, Mao C, Duraki D, Kim JE, Huang R, Helferich WG, Nelson ER, Park BH, Shapiro DJ: Estrogen-independent Myc overexpression confers endocrine therapy resistance on breast cancer cells expressing ERαY537S and ERαD538G mutations. Cancer Lett 2019, 442:373–382.

42. Wu T, Hu E, Xu S, Chen M, Guo P, Dai Z, Feng T, Zhou L, Tang W, Zhan L, et al: clusterProfiler 4.0: A universal enrichment tool for interpreting omics data. The Innovation 2021, 2:100141.

43. Kanehisa M: The KEGG database. Novartis Found Symp 2002, 247:91–101; discussion 101-103, 119-128, 244-152.

44. Kanehisa M, Furumichi M, Sato Y, Kawashima M, Ishiguro-Watanabe M: KEGG for taxonomy-based analysis of pathways and genomes. Nucleic Acids Res 2023, 51:D587–d592.

45. Liberzon A, Birger C, Thorvaldsdóttir H, Ghandi M, Mesirov JP, Tamayo P: The Molecular Signatures Database (MSigDB) hallmark gene set collection. Cell Syst 2015, 1:417–425.

46. Tickle T, Tirosh I, Georgescu C, Brown M, Haas B: inferCNV of the Trinity CTAT Project. 2019.

47. Kurtenbach S, Cruz AM, Rodriguez DA, Durante MA, Harbour JW: Uphyloplot2: visualizing phylogenetic trees from single-cell RNA-seq data. BMC Genomics 2021, 22:419.

48. Saelens W, Cannoodt R, Todorov H, Saeys Y: A comparison of single-cell trajectory inference methods. Nat Biotechnol 2019, 37:547–554.

49. Gamper J, Alemi Koohbanani N, Benet K, Khuram A, Rajpoot N: PanNuke: An Open Pan-Cancer Histology Dataset for Nuclei Instance Segmentation and Classification. In Digital Pathology; 2019//; Cham. Edited by Reyes-Aldasoro CC, Janowczyk A, Veta M, Bankhead P, Sirinukunwattana K. Springer International Publishing; 2019: 11–19.

50. Jin S, Guerrero-Juarez CF, Zhang L, Chang I, Ramos R, Kuan CH, Myung P, Plikus MV, Nie Q: Inference and analysis of cell-cell communication using CellChat. Nat Commun 2021, 12:1088.

51. Hänzelmann S, Castelo R, Guinney J: GSVA: gene set variation analysis for microarray and RNA-seq data. BMC Bioinformatics 2013, 14:7.

52. Banerji S, Cibulskis K, Rangel-Escareno C, Brown KK, Carter SL, Frederick AM, Lawrence MS, Sivachenko AY, Sougnez C, Zou L, et al: Sequence analysis of mutations and translocations across breast cancer subtypes. Nature 2012, 486:405–409.

53. Zhang Y, Cai Q, Shu XO, Gao YT, Li C, Zheng W, Long J: Whole-Exome Sequencing Identifies Novel Somatic Mutations in Chinese Breast Cancer Patients. J Mol Genet Med 2015, 9.

54. Wei W, Zhang X, Sun S, Xia B, Liang X, Cui Y, Gao S, Pang D: Assessment of basal-like breast cancer by circulating tumor DNA analysis. Oncol Lett 2018, 15:7389–7396.

55. Cho MS, Park CH, Lee S, Park HS: Clinicopathological parameters for circulating tumor DNA shedding in surgically resected non-small cell lung cancer with EGFR or KRAS mutation. PLoS One 2020, 15:e0230622.

56. Cailleux F, Agostinetto E, Lambertini M, Rothé F, Wu HT, Balcioglu M, Kalashnikova E, Vincent D, Viglietti G, Gombos A, et al: Circulating Tumor DNA After Neoadjuvant Chemotherapy in Breast Cancer Is Associated With Disease Relapse. JCO Precis Oncol 2022, 6:e2200148.

57. Chen Y, Pal B, Lindeman GJ, Visvader JE, Smyth GK: R code and downstream analysis objects for the scRNA-seq atlas of normal and tumorigenic human breast tissue. Scientific Data 2022, 9:96.

58. Jie X-L, Wei J-C, Wang D, Zhang X-W, Lv M-Y, Lin Y-F, Tan Y-S, Wang Z, Alifu A, Ji L, et al: CDC34 suppresses macrophage phagocytic activity and predicts poor response to immune checkpoint inhibitor in cancers. Cancer Letters 2025, 628:217822.

59. Bharde A, Nadagouda S, Dongare M, Hariramani K, Basavalingegowda M, Haldar S, D’Souza A, Jadhav B, Prajapati S, Jadhav V, et al: ctDNA-based liquid biopsy reveals wider mutational profile with therapy resistance and metastasis susceptibility signatures in early-stage breast cancer patients. J Liq Biopsy 2025, 7:100284.

60. Lo YMD, Han DSC, Jiang P, Chiu RWK: Epigenetics, fragmentomics, and topology of cell-free DNA in liquid biopsies. Science 2021, 372.

61. Dustin D, Gu G, Fuqua SAW: ESR1 mutations in breast cancer. Cancer 2019, 125:3714–3728.

62. Will M, Liang J, Metcalfe C, Chandarlapaty S: Therapeutic resistance to anti-oestrogen therapy in breast cancer. Nat Rev Cancer 2023, 23:673–685.

63. Takeshita T, Yamamoto Y, Yamamoto-Ibusuki M, Tomiguchi M, Sueta A, Murakami K, Omoto Y, Iwase H: Comparison of ESR1 Mutations in Tumor Tissue and Matched Plasma Samples from Metastatic Breast Cancer Patients. Transl Oncol 2017, 10:766–771.

64. Reinhardt K, Stückrath K, Hartung C, Kaufhold S, Uleer C, Hanf V, Lantzsch T, Peschel S, John J, Pöhler M, et al: PIK3CA-mutations in breast cancer. Breast Cancer Res Treat 2022, 196:483–493.

65. Bader AG, Kang S, Vogt PK: Cancer-specific mutations in PIK3CA are oncogenic in vivo. Proc Natl Acad Sci U S A 2006, 103:1475–1479.

66. Jenkins ML, Ranga-Prasad H, Parson MAH, Harris NJ, Rathinaswamy MK, Burke JE: Oncogenic mutations of PIK3CA lead to increased membrane recruitment driven by reorientation of the ABD, p85 and C-terminus. Nat Commun 2023, 14:181.

67. Zhang M, Zhou K, Wang Z, Liu T, Stevens LE, Lynce F, Chen WY, Peng S, Xie Y, Zhai D, et al: A Subpopulation of Luminal Progenitors Secretes Pleiotrophin to Promote Angiogenesis and Metastasis in Inflammatory Breast Cancer. Cancer Res 2024, 84:1781–1798.

68. Badve SS, Gökmen-Polar Y: Ductal carcinoma in situ of breast: update 2019. Pathology 2019, 51:563–569.

69. Mohan T, Deng L, Wang BZ: CCL28 chemokine: An anchoring point bridging innate and adaptive immunity. Int Immunopharmacol 2017, 51:165–170.

70. Bhati R, Patterson C, Livasy CA, Fan C, Ketelsen D, Hu Z, Reynolds E, Tanner C, Moore DT, Gabrielli F, et al: Molecular characterization of human breast tumor vascular cells. Am J Pathol 2008, 172:1381–1390.

71. Varadan V, Kamalakaran S, Gilmore H, Banerjee N, Janevski A, Miskimen KL, Williams N, Basavanhalli A, Madabhushi A, Lezon-Geyda K, et al: Brief-exposure to preoperative bevacizumab reveals a TGF-β signature predictive of response in HER2-negative breast cancers. Int J Cancer 2016, 138:747–757.

72. Sparano JA, Gray RJ, Makower DF, Pritchard KI, Albain KS, Hayes DF, Geyer CE, Jr., Dees EC, Perez EA, Olson JA, Jr., et al: Prospective Validation of a 21-Gene Expression Assay in Breast Cancer. N Engl J Med 2015, 373:2005–2014.

73. Cardoso F, Van’t Veer L, Rutgers E, Loi S, Mook S, Piccart-Gebhart MJ: Clinical application of the 70-gene profile: the MINDACT trial. J Clin Oncol 2008, 26:729–735.

74. Loi S, Johnston S, Arteaga C, Graff S, Chandarlapaty S, Goetz M, Desmedt C, Reis-Filho J, Sasano H, Rodrik-Outmezguine V, et al: Abstract PS06-01: Results from a pilot study exploring ctDNA detection using a tumor-informed assay in the monarchE trial of adjuvant abemaciclib with endocrine therapy in HR+, HER2-, node-positive, high-risk early breast cancer. Cancer Research 2024, 84:PS06-01–PS06-01.

75. Martin-Alonso C, Tabrizi S, Xiong K, Blewett T, Sridhar S, Crnjac A, Patel S, An Z, Bekdemir A, Shea D, et al: Priming agents transiently reduce the clearance of cell-free DNA to improve liquid biopsies. Science 2024, 383:eadf2341.

76. Li S, Zeng W, Ni X, Liu Q, Li W, Stackpole ML, Zhou Y, Gower A, Krysan K, Ahuja P, et al: Comprehensive tissue deconvolution of cell-free DNA by deep learning for disease diagnosis and monitoring. Proc Natl Acad Sci U S A 2023, 120:e2305236120.

77. Wang S, Li M, Zhang J, Xing P, Wu M, Meng F, Jiang F, Wang J, Bao H, Huang J, et al: Circulating tumor DNA integrating tissue clonality detects minimal residual disease in resectable non-small-cell lung cancer. J Hematol Oncol 2022, 15:137.

78. Hu Z, Chen H, Long Y, Li P, Gu Y: The main sources of circulating cell-free DNA: Apoptosis, necrosis and active secretion. Crit Rev Oncol Hematol 2021, 157:103166.

79. Kamada T, Togashi Y, Tay C, Ha D, Sasaki A, Nakamura Y, Sato E, Fukuoka S, Tada Y, Tanaka A, et al: PD-1(+) regulatory T cells amplified by PD-1 blockade promote hyperprogression of cancer. Proc Natl Acad Sci U S A 2019, 116:9999–10008.

80. Heldin CH, Rubin K, Pietras K, Ostman A: High interstitial fluid pressure - an obstacle in cancer therapy. Nat Rev Cancer 2004, 4:806–813.

81. Teixeira MF, Fahmy N, Kasi PM: Utility of Circulating Tumor DNA-Based Liquid Biopsies in Patients with Cancer Receiving Immunotherapy. Surg Oncol Clin N Am 2026, 35:399–414.

82. Fallah J, Ganguly S, Rayman P, Wei W, Balyimez A, Sitalaximi T, Lamenza M, Stephans KL, Dann P, Tendulkar RD, et al: Association of cell-free DNA (cfDNA) levels with myeloid-derived suppressor cells (MDSC) levels in blood of patients (pts) with muscle invasive (MI) and metastatic (met) bladder cancer (BC). Journal of Clinical Oncology 2019.

83. Steimle AK, Cho SY, Menon N, Jeraj R, Birbrair A, Albertini MR, Ma VT: ctDNA Dynamics Identifies Pseudoprogression in a Metastatic Melanoma Patient Treated With Nivolumab/Relatlimab. J Immunother 2025, 48:325–328.

84. Katsuno Y, Meyer DS, Zhang Z, Shokat KM, Akhurst RJ, Miyazono K, Derynck R: Chronic TGF-β exposure drives stabilized EMT, tumor stemness, and cancer drug resistance with vulnerability to bitopic mTOR inhibition. Sci Signal 2019, 12.

85. Sanz-Garcia E, Zhao E, Bratman SV, Siu LL: Monitoring and adapting cancer treatment using circulating tumor DNA kinetics: Current research, opportunities, and challenges. Sci Adv 2022, 8:eabi8618.

