## Supplementary Figures for "ctDNA Release Heterogeneity in Luminal Breast Cancer Revealed by Single-Cell Multi-Omics Spatial Analysis"

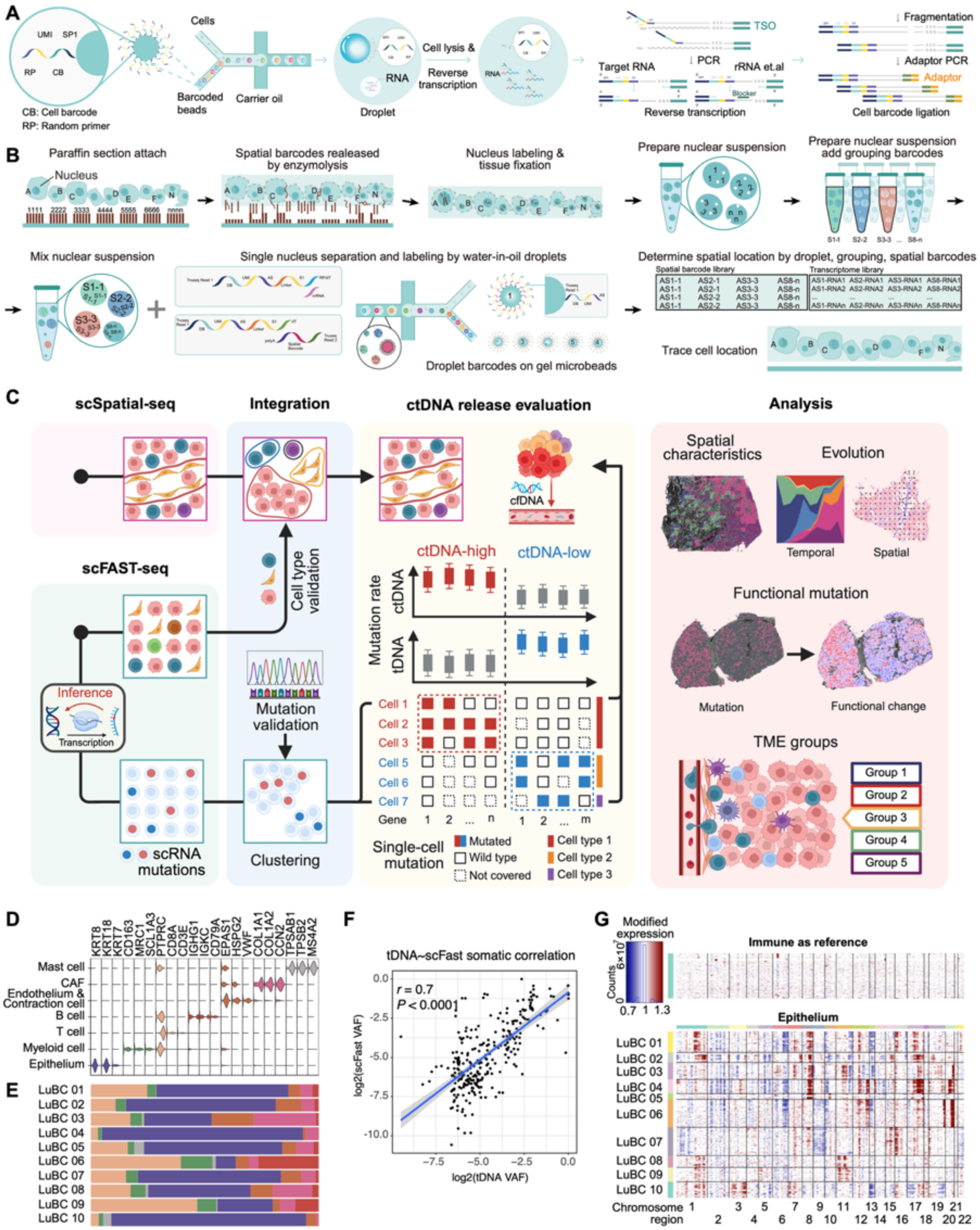


**Extended Data Fig. 1: Analytical workflows and technical validation.**

(*A*) Schematic illustration of the scFAST-seq library preparation protocol. (*B*) Schematic illustration of the scSpatial-seq experimental procedure. (*C*) Comprehensive workflow for the integration and multi-omic analysis of single-cell and spatial datasets. (*D*) Cell-type-specific marker gene expression across endothelial, immune, and epithelial cells. (*E*) Relative proportions of cellular components across all sequenced samples. (*F*) Correlation between variant allele frequencies (VAF) derived from scFAST-seq and bulk tissue DNA (tDNA; Pearson's $r = 0.70$, $P < 0.0001$).

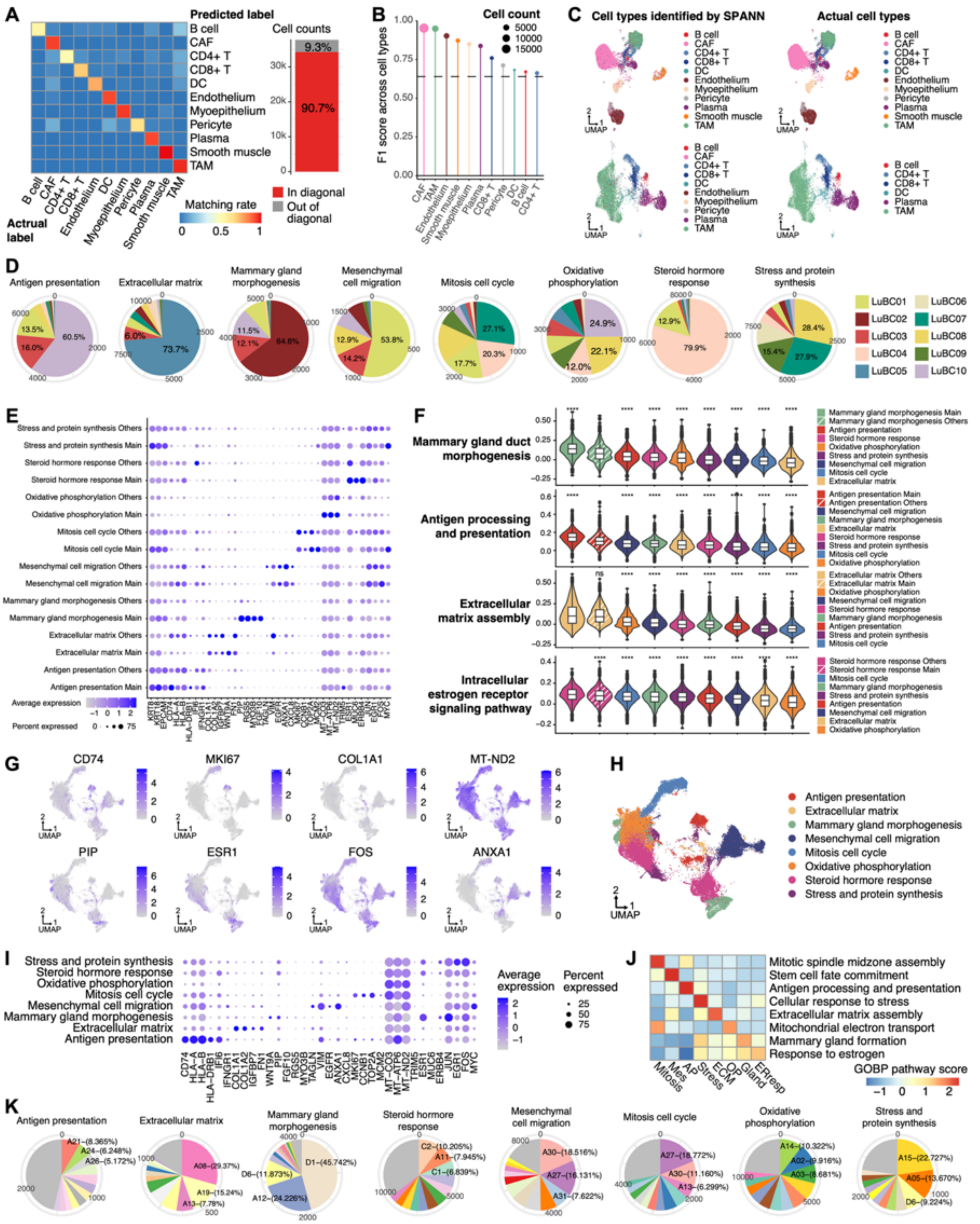


Extended Data Fig. 2: Validation of transferred cell-type annotation and cross-cohort robustness of epithelial states.

(*A*) Confusion matrix showing the consistency between transferred labels and manual annotations, with the diagonal indicating the proportion of correctly assigned cells. (*B*) F1 scores evaluating the precision and recall of label transfer across various immune cell subtypes. (*C*) UMAP visualizations comparing SPANN-identified cell types with manual annotations for non-epithelial cells (top) and immune cell populations (bottom). (*D*) Relative distribution of individual patients across the eight malignant epithelial states. (*E*) Expression of marker genes in the total cell population versus major patient-depleted fractions. (*F*) Functional enrichment scores of Gene Ontology Biological Process (GOBP) pathways across the eight epithelial states and major patient-depleted fractions. (*G*) UMAP visualization of core marker genes in the external validation cohort (BC50). (*H*) UMAP projection of BC50 epithelial cells classified according to the eight-state framework. (*I*) Expression profiles of signature genes defining the eight epithelial states in the BC50 cohort. (*J*) GOBP pathway enrichment scores for the eight epithelial states within the BC50 cohort. (*K*) Relative distribution of individual patients across the eight epithelial states in the BC50 cohort.

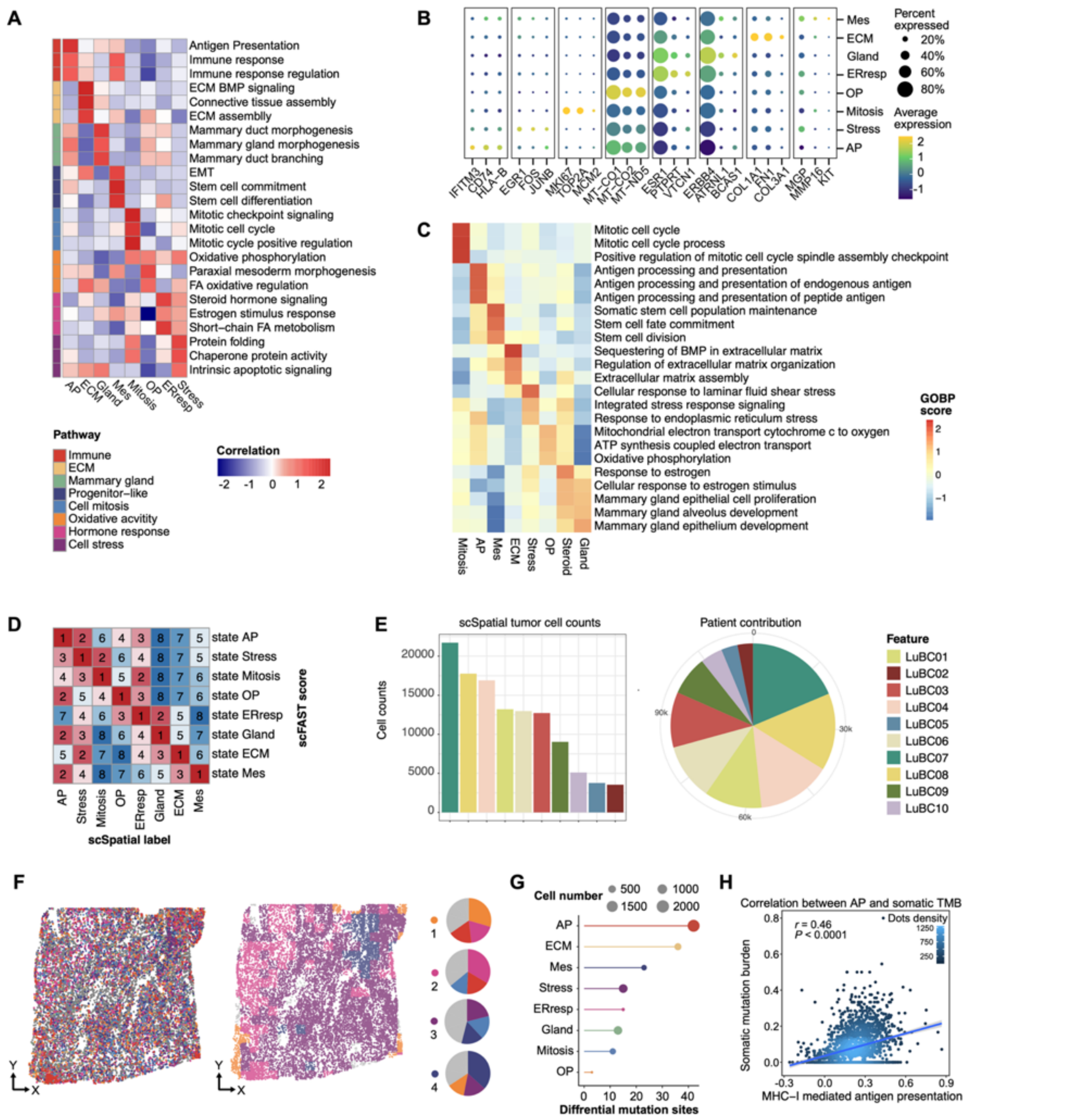


**Extended Data Fig. 3: Label transfer and spatial distribution of epithelial states.**

(*A*) Representative biological functions of the eight epithelial states in the scFAST-seq dataset. (*B*) Dot plot illustrating the expression of signature genes defining each epithelial state. (*C*) Heatmap of representative GOBP pathway scores for the eight epithelial states in the scSpatial-seq dataset. (*D*) Rank-based assignment matrix showing the concordance between scFAST-seq scores (rows) and scSpatial-seq labels (columns). Numeric values (1–8) indicate the rank of scFAST-seq scores for each spatial label. (*E*) Cell counts and relative distribution of individual patients within the scSpatial-seq cohort. (*F*) Spatial mapping of epithelial states and regions in a representative sample (left), with pie charts indicating the cellular compositions of major epithelial regions (right). (*G*) Dot plots illustrating the number of differential mutation sites and mutated cell counts across the eight states. (*H*) Positive correlation between MHC-I mediated antigen presentation scores and somatic mutation burden (Pearson's $r = 0.46$, $P < 0.0001$).

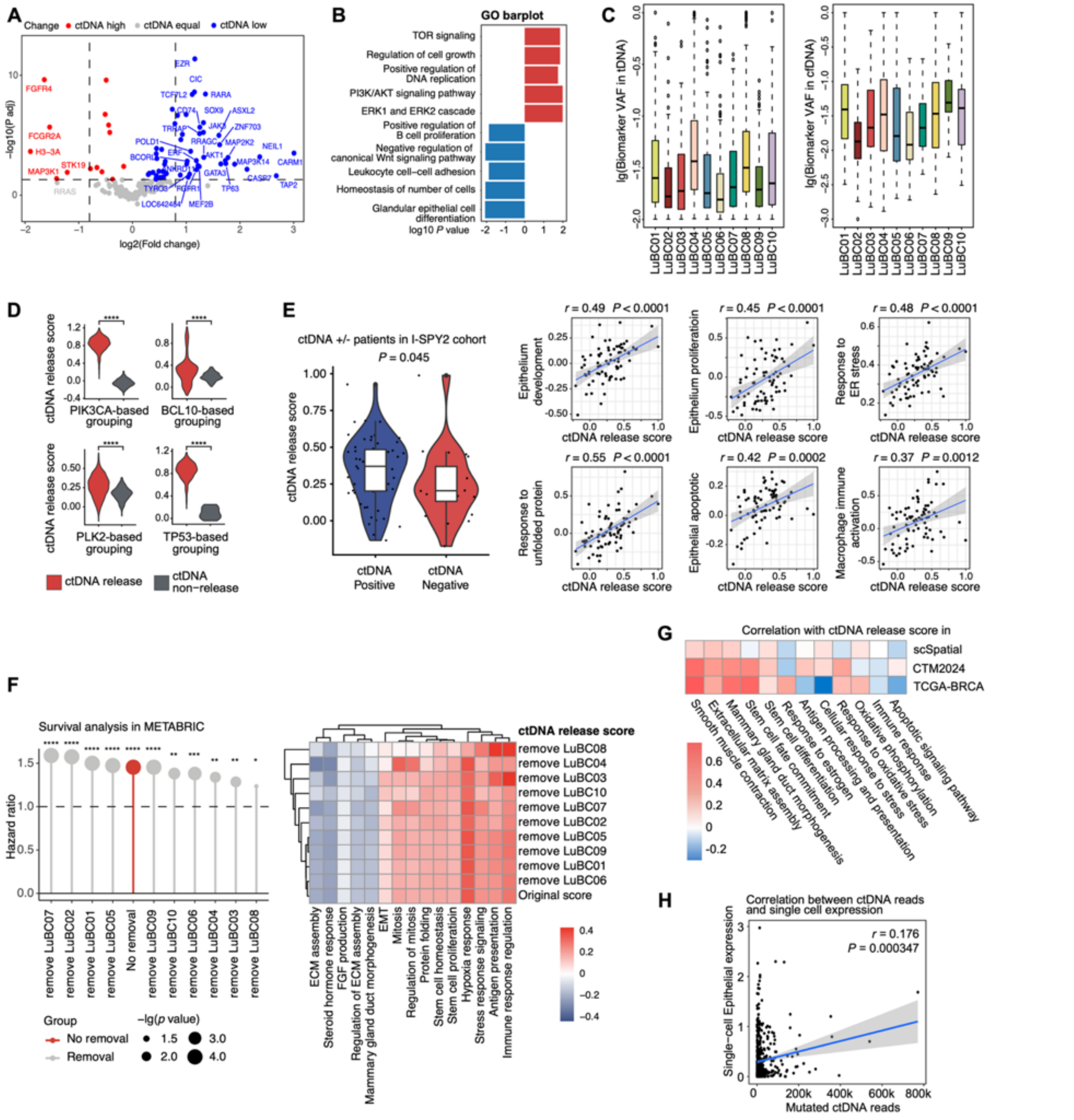


**Extended Data Fig. 4: Identification of ctDNA biomarkers and multi-cohort validation of the ctDNA release score.**

(*A*) Volcano plot comparing cfDNA and tDNA variant allele frequencies (VAFs) across ten patients. (*B*) GO enrichment of functional pathways for cfDNA-high versus cfDNA-low biomarkers. (*C*) VAF distributions of biomarker mutations in cfDNA and tDNA. (*D*) Comparison of ctDNA release scores in epithelial cells, stratified by the presence or absence of specific gene mutations in ctDNA. (*E*) Validation in the I-SPY2 cohort: Higher ctDNA release scores in ctDNA-positive versus ctDNA-negative patients at baseline (left), and their correlation with functional signatures (right; Pearson's). (*F*) Leave-one-out sensitivity analysis: 5-year overall survival (OS) in METABRIC stratified by ctDNA release scores (left; Hazard ratio and log-rank *P* shown), and correlation between ctDNA release scores and functional scores (right). (*G*) Correlation between ctDNA release scores and representative biological functions across multiple external datasets. (*H*) Positive correlation between single-cell epithelial expression and mutated ctDNA reads in epithelial cells (Pearson's $r = 0.176$, $P = 0.000347$).

**Statistics:** *P* values determined by two-tailed Student's *t*-test (*D, E*) or log-rank test (*F*).

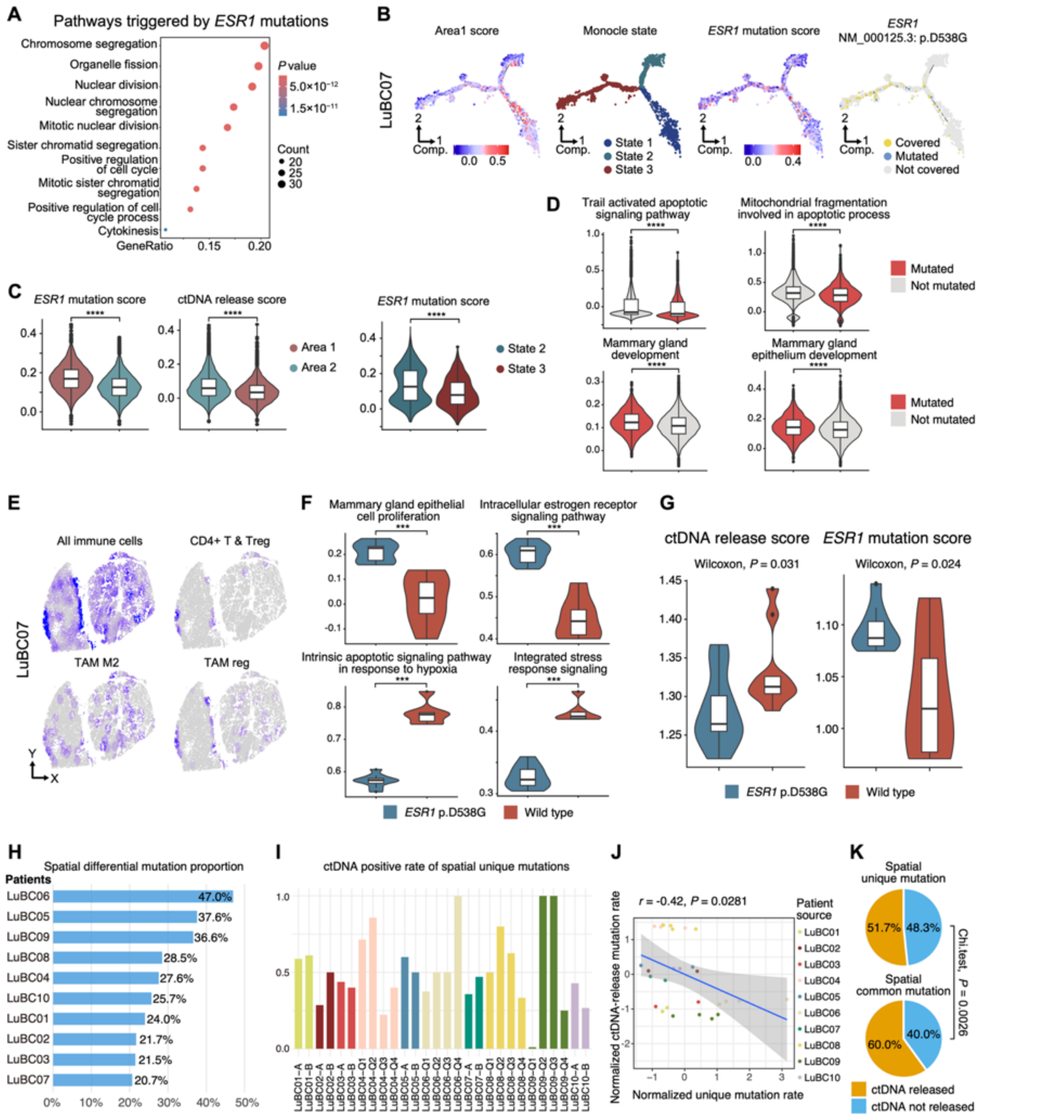


**Extended Data Fig. 5: *ESR1* mutations and spatial heterogeneity associated with ctDNA release.**

(*A*) Dot plot showing the gene ratio of *ESR1*-related pathways triggered by *ESR1* mutations. (*B*) Cellular trajectories of Area 1 score, Monocle state, *ESR1* mutation score, and *ESR1* mutation state in a representative scFAST-seq sample. (*C*) Violin plots comparing *ESR1* mutation and ctDNA release scores between Area 1 and Area 2 (left), and *ESR1* mutation scores between State 2 and State 3 cells (right). (*D*) Differential pathway expression between inferred mutated and non-mutated cells in scSpatial-seq data. (*E*) Spatial distribution of immune cell subsets (CD4+ T/Treg, TAM M2, and TAM reg) in a representative sample. (*F–G*) External validation of *ESR1* D538G: Differential functional pathways (*F*) and comparison of ctDNA release and *ESR1* mutation scores (*G*) between wild-type and *ESR1* D538G mutant cells. (*H*) Proportion of mutations exhibiting significant spatial heterogeneity across patients (Chi-square test, $P < 0.05$). (*I*) Detection rate in ctDNA for spatial unique mutations across all regions. (*J*) Correlation between the proportion of region-specific unique mutations and their corresponding ctDNA detection rates (Pearson's $r = 0.42$, $P = 0.0281$). (*K*) Comparison of ctDNA detection rates between spatial unique and common mutations.

**Statistics:** *P* values determined by two-tailed Student's *t*-test (*A, C, D, F*) or Wilcoxon test (*E*) with BH correction; **** $P < 0.0001$.

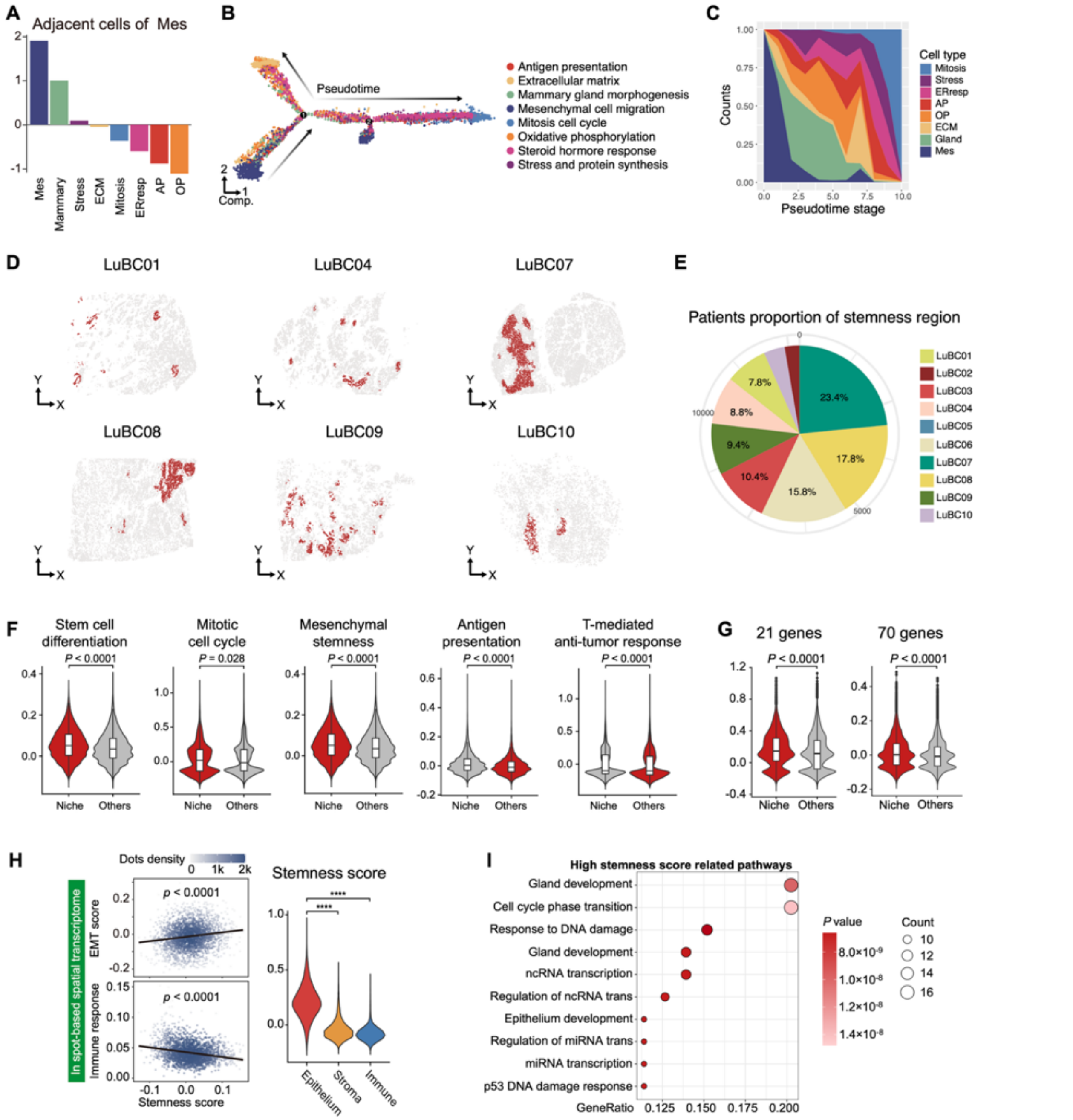


**Extended Data Fig. 6: Spatial organization, differentiation trajectories, and characterization of stemness regions.**

(*A*) Standardized proportions of adjacent cell types surrounding the Mesenchymal (Mes) state identified by spatial adjacency analysis. (*B–C*) Trajectory analysis of the total cell population (*B*) and the relative proportion of epithelial states across pseudotime (*C*). (*D*) Spatial visualization of stemness regions across multiple samples (LuBC01, 04, 07, 08, 09, and 10). (*E*) Relative cellular contribution of individual patients within stemness regions. (*F–G*) Violin plots comparing functional signatures—including stem cell differentiation, mitotic cell cycle, mesenchymal stemness, antigen presentation, T-mediated anti-tumor response, and 21-gene/70-gene scores—between epithelial cells within and outside stemness regions. (*H*) Correlation analysis of stemness scores with EMT and immune response scores (left), and comparison of stemness scores across three distinct cell types (right). (*I*) Dot plot showing the gene ratio of functional pathways enriched in cells with high stemness scores.

**Statistics:** *P* values determined by two-tailed Student's *t*-test with BH correction (*F, G, H, I*); correlations in (*H*) identified by Pearson analysis ($r = 0.322$, $P < 0.0001$ for EMT; $r = -0.380$, $P < 0.0001$ for immune response).

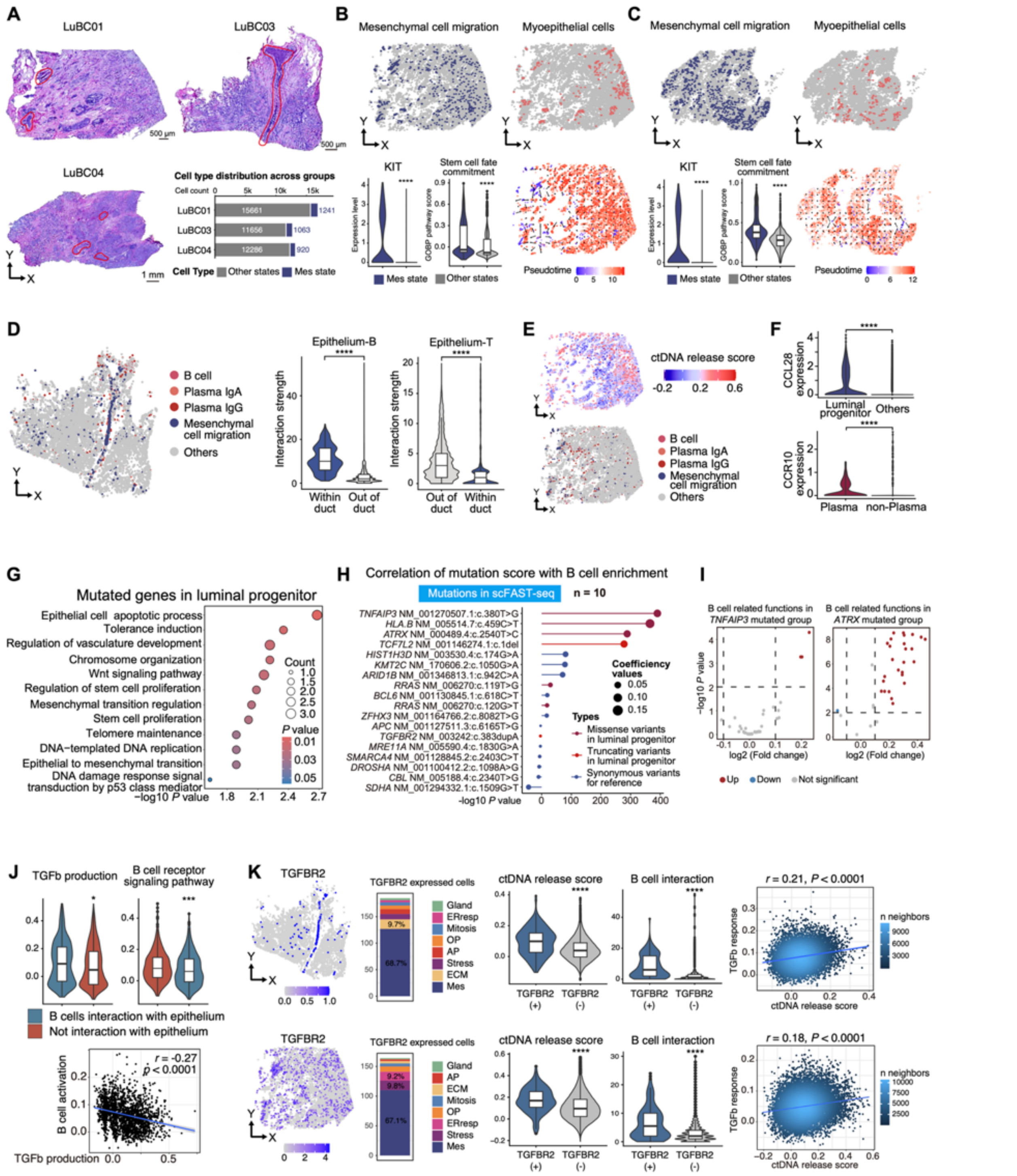


**Extended Data Fig. 7: Spatial characterization of ductal carcinoma and immune-epithelial interactions modulating ctDNA release.**

(*A*) Pathological annotation of mammary ducts and ductal carcinoma in situ (DCIS) in LuBC01, 03, and 04 tissue sections, with mesenchymal cell migration (Mes) state distribution. (*B–C*) scSpatial profiling and spatial trajectory analysis of DCIS regions in patients LuBC01 and LuBC04. (*D*) Immune interactions in LuBC03, showing enhanced B cell and diminished T cell interactions within mammary ducts. (*E*) Spatial distribution of the Mes state, B cells, and ctDNA release scores in LuBC01. (*F*) Violin plots showing CCL28 expression in Mes versus other states (top) and CCR10 expression in plasma versus non-plasma (bottom). (*G*) Enriched biological pathways associated with mutated genes in luminal progenitor cells. (*H*) B cell enrichment across eighteen mutation scores, categorized by missense/truncating variants and synonymous reference. (*I*) Comparison of B cell-related functions in patients with or without *TNFAIP3* (left) and *ATRX* (right) mutations in the TCGA dataset. (*J*) Functional characterization of B cells with epithelial interactions, showing increased TGFB production and decreased BCR signaling; TGFB production negatively correlates with B cell activation. (*K*) Comprehensive analysis on TGFBR2, including spatial featureplots, cell proportions, and associations with B interactions and ctDNA release scores in LuBC01 and LuBC03.

**Statistics:** *P* values determined by two-tailed Student's *t*-test with BH correction; * $P < 0.05$; ** $P < 0.01$; *** $P < 0.001$; **** $P < 0.0001$.

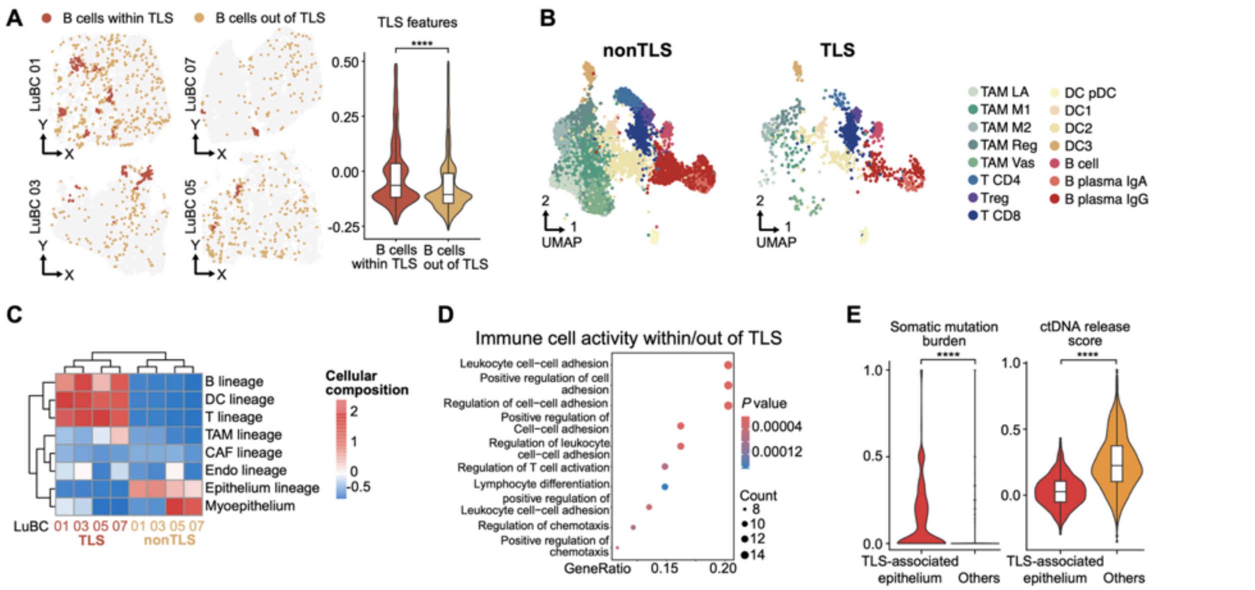


**Extended Data Fig. 8: Spatial characterization of TLS and its association with ctDNA release.**

(*A*) Illustrations of B-cell spatial visualization (left) and TLS features (right) in TLS and non-TLS regions across four representative samples. (*B*) UMAP projection of immune cell subtypes of immune cells within and without TLS in scFAST-seq data. (*C*) Cellular lineage differences between immune cells within and without TLS in four samples. (*D*) Dot plot depicting enriched biological pathways in mutated genes in the luminal progenitor. (*E*) Violin plots showing somatic mutation burden and ctDNA release scores in TLS-associated epithelial cells.

**Statistics:** *P* values determined by two-tailed Student's *t*-test with BH correction (*A, E*); **** $P < 0.0001$.

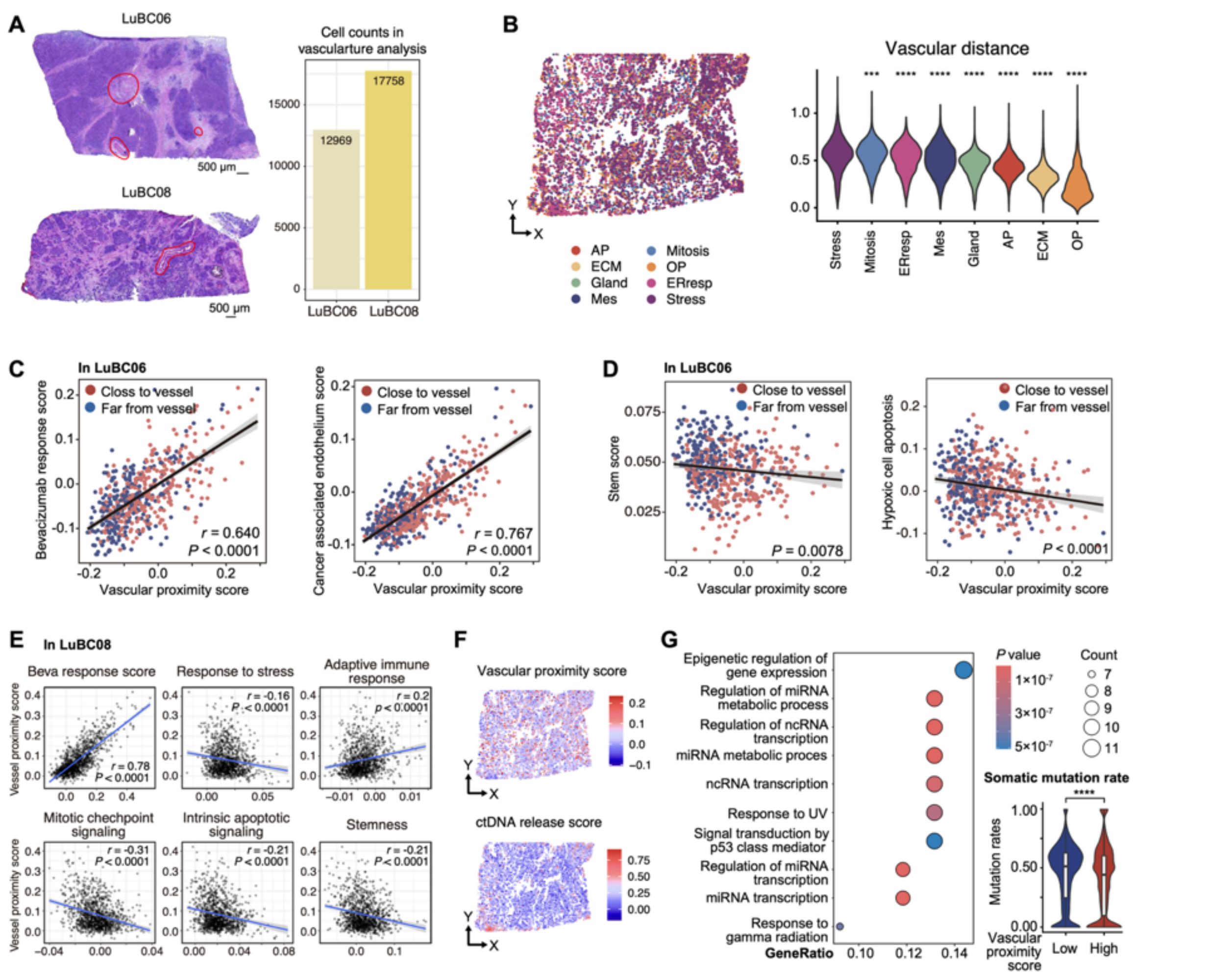


**Extended Data Fig. 9: Label transfer and spatial distribution of epithelial states.**

(*A*) Representative tissue sections from LuBC06 and LuBC08 showing vascular annotation (left) and corresponding scSpatial cell counts (right). (*B*) Spatial DimPlot and violin plots depicting vascular distance across eight epithelial states in LuBC08. (*C*) Correlation analyses in LuBC06 showing positive associations between vascular proximity score and bevacizumab response score, and cancer-associated endothelium score. (*D*) Correlation analyses in LuBC06 showing vascular proximity score correlates negatively with the stemness score and hypoxic cell apoptosis score. (*E*) Correlations in

LuBC08 showing associations between vascular proximity score and various functional scores, including bevacizumab response, stress response, adaptive immune response, mitotic checkpoint signaling, intrinsic apoptotic signaling, and stemness. (*F*) Spatial featureplots in LuBC08 illustrating the relationship between vascular proximity score and ctDNA release score. (*G*) Dot plot depicting enriched biological pathway of differentially mutated genes between epithelial cells with high or low vascular proximity scores.

**Statistics**: *P* values determined by two-tailed Student's *t*-test with BH correction (*B, G*) or Pearson's correlation (*C–E*); *** $P < 0.001$, **** $P < 0.0001$.

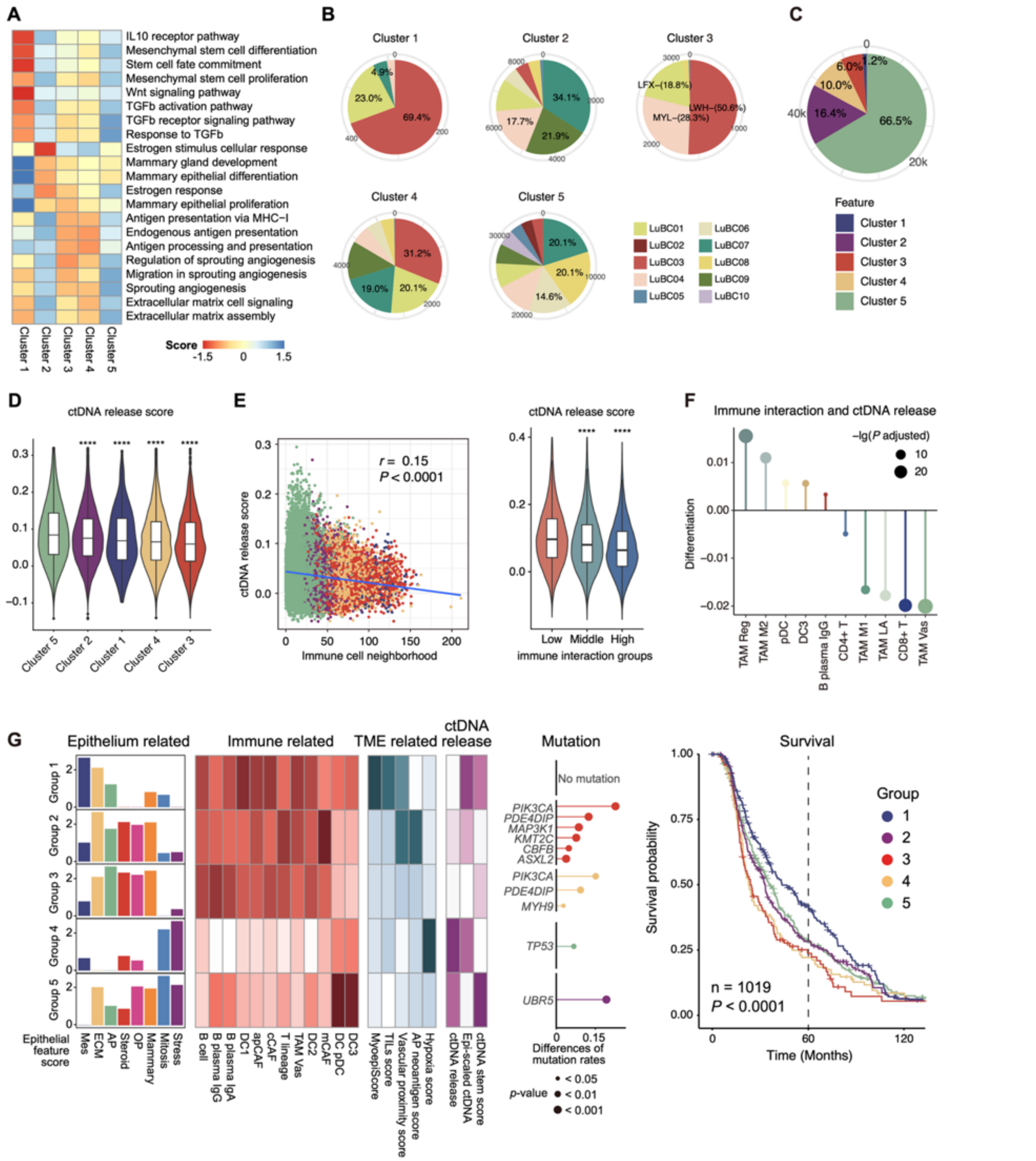


**Extended Data Fig. 10: Immune interaction landscapes and their association with ctDNA release.**

(*A*) Heatmap showing the functional characteristics of five epithelial clusters. (*B*) Pie plots showing the proportional distribution of patients across each TME neighborhood cluster. (*C*) Pie chart illustrating the proportion of the five clusters among all scSpatial malignant cells. (*D*) Violin plots displaying ctDNA release scores for the five clusters, ordered from highest to lowest median scores. (*E*) Left: Scatter plot showing the correlation between total immune neighborhood counts and ctDNA release scores. Right: Violin plots comparing ctDNA release scores across epithelial cells stratified by tertiles of immune neighborhood counts. (*F*) Lolipop plots comparing ctDNA release scores between epithelial cells that interact with specific immune cell subsets and those that do not exhibit such interactions. (*G*) Patient risk stratification and characteristics in the TCGA dataset, including eight epithelial features, immune-related scores, TME-related scores, ctDNA release scores, differential mutation genes, and Kaplan-Meier survival curves for five patient groups.

**Statistics**: *P* values determined by two-tailed Wilcoxon rank-sum tests with Benjamini–Hochberg correction (*D, E*), Pearson's correlation (*E*), or log-rank test (*G*); * $P < 0.05$; ** $P < 0.01$; *** $P < 0.001$; **** $P < 0.0001$.
